# Multivariate analysis of metabolic state vulnerabilities across diverse cancer contexts reveals synthetically lethal associations

**DOI:** 10.1101/2023.11.28.569098

**Authors:** Cara Abecunas, Audrey D. Kidd, Ying Jiang, Hui Zong, Mohammad Fallahi-Sichani

**Author notes:** Present address: Novartis Institutes for BioMedical Research, Cambridge, MA 02139.

## Abstract

Targeting the distinct metabolic needs of tumor cells has recently emerged as a promising strategy for cancer therapy. The heterogeneous, context-dependent nature of cancer cell metabolism, however, poses challenges in identifying effective therapeutic interventions. Here, we utilize various unsupervised and supervised multivariate modeling approaches to systematically pinpoint recurrent metabolic states within hundreds of cancer cell lines, elucidate their association with tumor lineage and growth environments, and uncover vulnerabilities linked to their metabolic states across diverse genetic and tissue contexts. We validate key findings via analysis of data from patient-derived tumors and pharmacological screens, and by performing new genetic and pharmacological experiments. Our analysis uncovers new synthetically lethal associations between the tumor metabolic state (e.g., oxidative phosphorylation), driver mutations (e.g., loss of tumor suppressor PTEN), and actionable biological targets (e.g., mitochondrial electron transport chain). Investigating the mechanisms underlying these relationships can inform the development of more precise and context-specific, metabolism-targeted cancer therapies.

## Introduction

Cancer cells rely on metabolic pathways for a variety of functions, including proliferation, survival, and migration, making metabolism an attractive anti-cancer therapeutic target ^1^. Identifying effective metabolic targets for cancer therapy, however, has proven challenging ^2^. Tumor cells exhibit a wide range of metabolic states that vary by their tissue of origin, developmental stage, and genetic alterations ^3^. They also have remarkable ability to rewire their intracellular metabolic networks in response to diverse environmental cues and changing metabolic demands ^4,5^. The context-dependent nature of cancer cell metabolism makes it difficult to identify unique metabolic pathways that are uniformly required for the diverse population of cancer cells across heterogeneous tumors but are not essential for healthy cells. Despite this challenge, the discovery of novel cancer-specific metabolic dependencies has provided promising opportunities via, for example, targeting genetically mutated enzymes whose upregulated activity is essential for tumor growth ^6,7^, or blocking metabolic pathways on which tumor cells with specific oncogenic alterations have developed extraordinarily high dependency ^8–10^. Uncovering such context-specific metabolic alterations and dependencies can guide new approaches to target cancer cells.

The search for novel anti-cancer metabolic targets has significantly benefited from recent developments in characterization of large panels of diverse cancer cell lines, including their genetic dependency maps, genomic, transcriptomic, and metabolomic features ^11,12^. Systematic studies have generated and integrated such large-scale datasets to characterize the landscape of metabolic pathway alterations and dependencies across cancer cell lines, demonstrating the context-specific nature of metabolic dependencies, and identifying new metabolic vulnerabilities in cancer cells ^13–18^. Building upon promising outcomes from these studies, a critical next step would be to develop computational models that reveal metabolic state-specific gene or pathway co-dependencies to discover potential synthetic lethalities imposed by the cancer genotype or tissue context ^2^. Because metabolism is heavily dependent on subcellular organelle functions and integrated with processes such as kinase signaling, transcription and chromatin modifications ^4,19–22^, it is likely that a multivariate approach may reveal novel co-dependencies among these processes and heterogeneous metabolic states. Importantly, such co-dependencies, which may not be initially apparent, could be exploited to guide the development of new drugs, or repurposing of existing ones, to target cancer cells more precisely.

In this study, we test the hypothesis that genetic, and tissue context-specific therapeutic targets associated with heterogeneous metabolic states in cancer cells could be identified through conditional synthetic lethality. We integrate transcriptomics data from the Cancer Cell Line Encyclopedia, spanning 41 major cancer types, with their mutation profiles and genome-wide DepMap gene dependency scores ^12^. We find heterogeneities in some metabolic pathways (such as linoleic acid, phenylalanine, ascorbate and aldarate metabolism) was highly associated with cancer type, while cell line-to-cell line variability in other pathways, such as oxidative phosphorylation (Oxphos), could not be explained by differences in cancer type or lineage. Furthermore, Oxphos was the most variable pathway across individual cell lines in at least 11 major cancer types with little association with their growth media composition or culture condition, suggesting that cell-intrinsic factors drive the majority of Oxphos heterogeneity in these cell lines. Therefore, we employ multivariate modeling to uncover Oxphos^High^ and Oxphos^Low^ state-specific vulnerabilities in cell lines representing these cancers. Furthermore, we find some specific driver mutations or tissue contexts significantly amplify the impact of Oxphos state-specific gene dependencies. Loss of PTEN, for example, predicts increased dependency on mitochondrial respiratory chain and enhanced sensitivity to pharmacological inhibitors of mitochondrial ATP synthase in Oxphos^High^ tumor cells. Our approach, therefore, provides a path to identify context-specific, metabolic state-dependent synthetic lethalities that could be exploited to guide more precise cancer therapeutic opportunities.

## Results

### Transcriptomics analysis of cancer cell lines reveals cancer type-associated heterogeneities in metabolic pathways

To systematically evaluate metabolic similarities and differences across cell lines representing diverse cancer types, we used RNA sequencing data from the Cancer Cell Line Encyclopedia (CCLE) ^23^. By including all cancer types of which 5 CCLE cell lines or more were available, we analyzed 1,341 cell lines spanning 41 major cancer types (Figure 1A). We used uniform manifold approximation and projection (UMAP) clustering to visualize the degree to which the expression of 1,620 genes representing 85 different metabolic pathways (obtained from the KEGG database ^24,25^) varied with cancer type (Figure 1B). The UMAP projection revealed clusters of cell lines that correlated with cancer types based on their metabolic gene expression. Examples of cancers that clustered most distinguishably by UMAP were cell lines representing acute myeloid leukemia, B-and T-lymphoblastic leukemia/lymphoma, diffuse glioma, Ewing sarcoma, Hodgkin and non-Hodgkin lymphomas, neuroblastoma, melanomas, liposarcoma, rhabdoid cancer, and myeloproliferative neoplasms (Figures 1B and S1A). UMAP also revealed patterns of association between cell lines representing distinctive tissue lineages. For example, cell lines of the hematopoietic lineage (including leukemias, lymphomas, and myeloproliferative neoplasms) were all clustered together and separated from non-hematopoietic cell lines (Figure 1B), and within the hematopoietic cancers, lymphoid and myeloid cell lines also clustered separately (Figure S1B). Clustering of melanoma cell lines of cutaneous and ocular subtype or cell lines of the neuroendocrine lineage (including small cell lung cancer and neuroblastoma) were additional examples, where metabolic gene expression signatures revealed similarities among cancer types with related lineages (Figures 1B and 1C). Interestingly, however, within some cancer types such as non-small cell lung cancer (NSCLC), ovarian epithelial tumor, or esophagogastric adenocarcinoma, we observed substantial differences among individual cell lines (Figures 1C and S1A). In the case of NSCLC, we asked whether these differences might be explained by the NSCLC subtype, including squamous cell carcinoma, adenocarcinoma, and large cell carcinoma. Cancer cell lines representing these subtypes also did not cluster together in the UMAP visualization (Figure S1C), suggesting that other oncogenic events or environmental factors might have a stronger effect on patterns of metabolic gene expression among these cancer cell lines.

**Figure 1.**
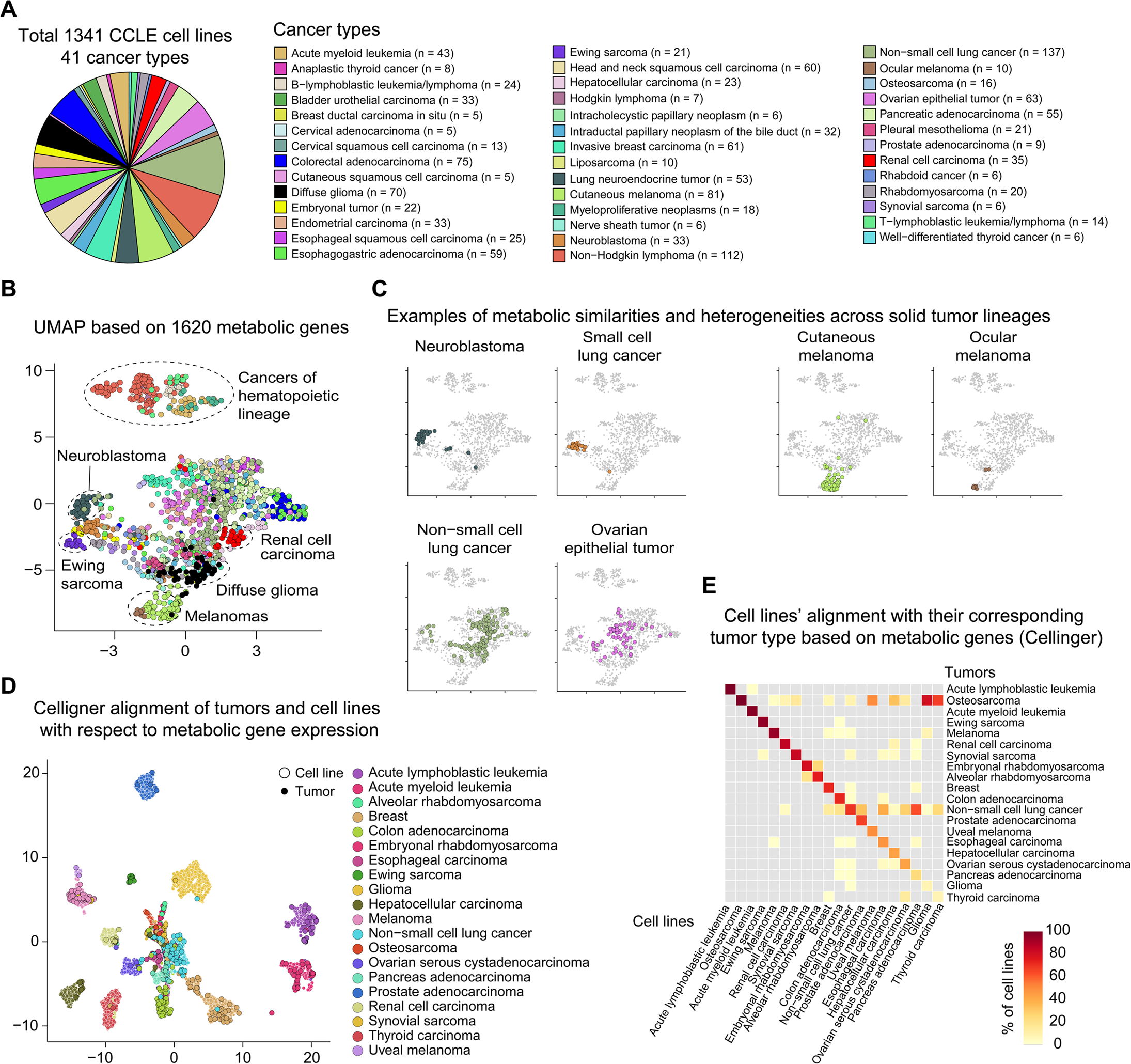
UMAP analysis of metabolic gene expression reveals clusters of cancer cell lines associated with tumor type and developmental lineage. (A) Total 1,341 CCLE cell lines, representing 41 cancer types, were included in the analysis. Number of cell lines for each cancer type are shown. (B) UMAP visualization of cell lines based on the expression levels of 1,620 metabolic genes. Cell lines are colored based on their cancer type (as shown in A). (C) Examples of metabolic similarities across tumor lineages, including neuroblastoma vs. small cell lung cancer (top left) and cutaneous vs. ocular melanoma (top right) or heterogeneities within non-small cell lung cancer and ovarian epithelial tumor cell lines (bottom) are shown. Individual cell lines from indicated cancer types are highlighted on UMAP plots. (D) UMAP visualization of Celligner-aligned cell lines and patient tumors based on the expression levels of 1,604 metabolic genes. Cell lines and tumors are colored based on their cancer type. (E) Percentage of cell lines of each cancer type that are grouped with their corresponding tumor type of origin using Celligner-aligned metabolic gene expression.

To test whether differences in metabolic gene expression across cell lines of diverse cancer types matched their tumor type of origin, we used Celligner ^26^, an unsupervised method to integrate and align RNA sequencing data from CCLE cell lines and patient-derived tumors included in TCGA, TARGET, and Treehouse databases ^27,28^. The Celligner-aligned metabolic gene expression profiles largely clustered together by cancer type in the UMAP projection (Figure 1D). On average, ∼66% of cell lines from each cancer type matched their tumor type of origin (Figure 1E). The alignment based on metabolic gene expression was reasonably high (>50%) for 14 out of 20 tested cancer types and particularly high (>90%) for leukemias, osteosarcoma, and Ewing sarcoma. The six cancer types showing <50% alignment between tumors and cell lines were esophageal carcinoma (48%), hepatocellular carcinoma (43%), ovarian serous cystadenocarcinoma (43%), pancreatic adenocarcinoma (24%), glioma (9%), and thyroid carcinoma (8%) (Figure 1E). Overall, these results reveal relationships between the expression of metabolic genes and cell lines of distinct cancer types, that are consistent with the tumor type of origin for most cancer types.

Next, we sought to perform a systematic statistical analysis to quantify these relationships at the level of individual metabolic pathways. To this end, we computed a score representing the activity of each of the 85 metabolic pathways for each cancer type relative to all cancer types using a three-step procedure adapted from Xiao *et al* ^29^ (see Methods). A metabolic pathway activity score <1 for a cancer type represents reduced pathway activity in that cancer type in comparison with the average pathway activity across all cancer types; scores >1 represent increased activity; and a score of 1 represents activity levels equivalent to the average over all cancer types. Among the 85 KEGG metabolic pathways, 73 exhibited significantly increased or decreased activity in at least one cancer type (Figure 2A). Hierarchical clustering revealed associations between pathway activity scores and each of the 41 cancer types. Cell lines from several haemopoietic cancers (e.g., B-and T-lymphoblastic leukemia/lymphoma, non-Hodgkin lymphoma, acute myeloid leukemia, and myeloproliferative neoplasms) exhibited overall reduced activity in a wide range of metabolic pathways in comparison with most other cancer types (Figures 2A and S2A). The activity of multiple metabolic pathways such as linoleic acid metabolism, metabolism by cytochrome P450, retinol metabolism, ascorbate and aldarate metabolism, and steroid hormone biosynthesis was substantially lower in these hematopoietic cancers relative to most other cancers. The activity of these metabolic pathways, however, were elevated in cell lines of non-small cell lung cancer, lung neuroendocrine tumor, hepatocellular and esophageal squamous cell carcinomas (Figure 2A).

**Figure 2.**
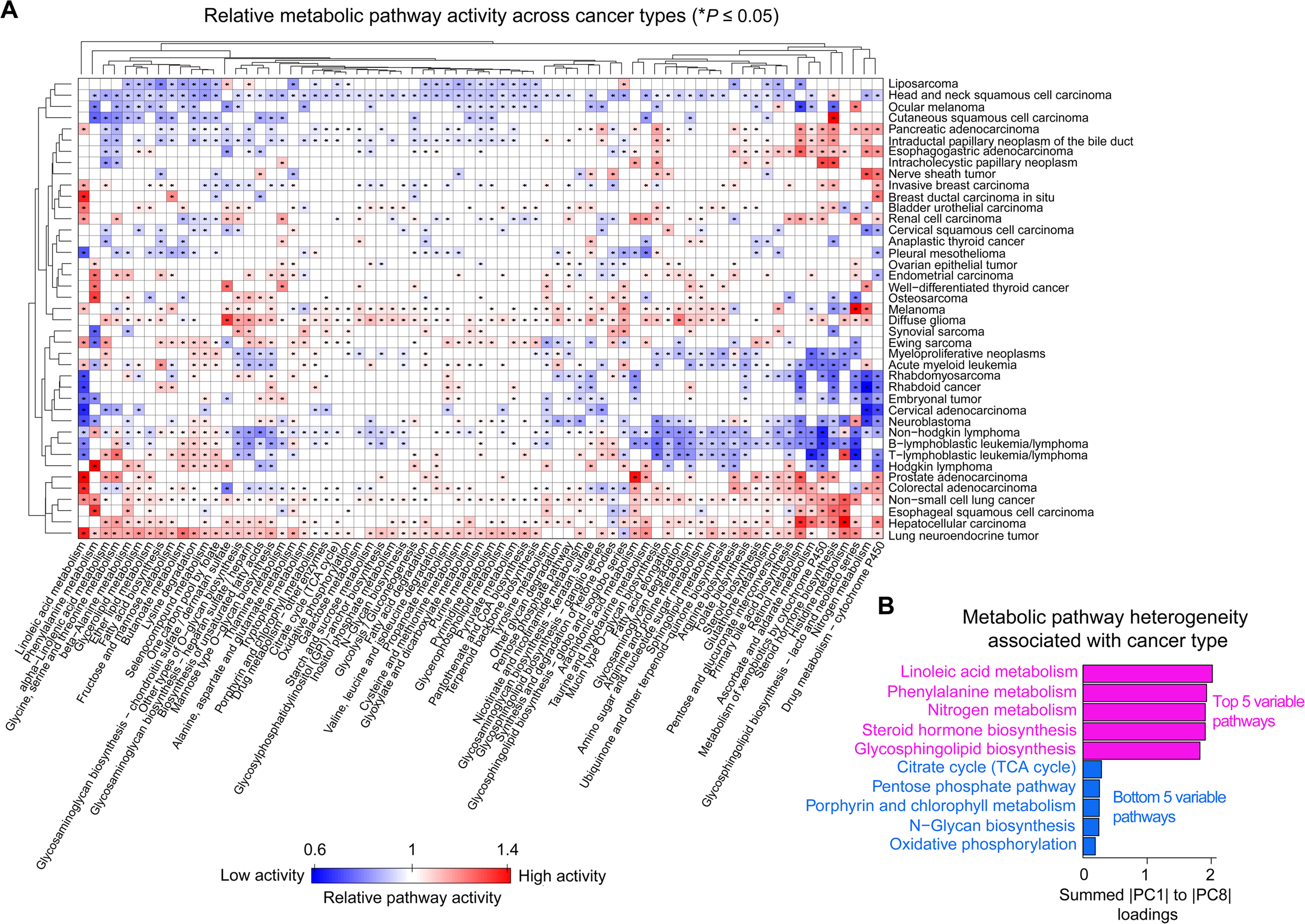
Metabolic pathway analysis of cancer cell lines reveals cancer type-associated variations in the activity of pathways. **(A)** Hierarchical clustering of relative activity scores for 73 significantly variable metabolic pathways across 41 cancer types. A metabolic pathway activity score < 1 for a cancer type represents reduced pathway activity in that cancer type in comparison with the average pathway activity across all cancer types; scores > 1 represent increased activity; and a score of 1 represents activity levels equivalent to the average over all cancer types. Pairs of pathway/cancer type with significant pathway activity (*P* ≤ 0.05) are highlighted with an asterisk (*). **(B)** Ranking of metabolic pathways based on the extent to which their heterogeneity is associated with cancer type, evaluated by computing the absolute sum of PCA loadings for each pathway over the first eight principal components capturing >80% of overall variance in data. The top 5 and bottom 5 variable pathways are shown.

To identify those metabolic pathways that contribute most profoundly to cancer type-associated variabilities, we performed principal component analysis (PCA) on pathway activity scores across 41 cancer types. The first eight principal components (PCs) captured >80% of the overall variance in data (Figure S2B). To quantify the relative impact of each pathway on data variance in the principal component space, we computed the absolute sum of PCA loadings for each pathway over the first eight PCs. This way of analysis would give a higher weight to metabolic pathways that are important in defining a larger number of principal components. Among the top 10 variable pathways across all cancer types were pathways involved in lipid, amino acid, and vitamin metabolism, such as linoleic acid, phenylalanine and histidine metabolism, steroid hormone, and glycosphingolipid biosynthesis, and ascorbate and aldarate metabolism (Figures 2B and S2C). On the other hand, the pathways with the least amount of overall variability across different cancer types were energy-producing and carbohydrate metabolism pathways, such as oxidative phosphorylation (Oxphos), citrate (TCA) cycle, pentose phosphate pathway, and N-glycan biosynthesis (Figures 2B and S2C). We repeated this analysis, focusing on PCs that captured either >70% or >90% of overall variance in data and reached similar conclusions (Figure S2D).

We then asked whether such cancer type-associated variability in metabolic pathways could be captured directly through metabolomics data analysis. UMAP analysis of data on 136 metabolites profiled for 875 CCLE cell lines ^11^ (representing 34 cancer types) revealed a clear separation between hematopoietic and solid-tissue cancers (Figure S2E), an observation that agreed with metabolic gene expression analysis. Using metabolite abundance alone, however, UMAP clustering did not reveal subtle cancer lineage-specific differences that were captured by the expression of 1,620 metabolic genes (Figure 1B). Nevertheless, focusing on the top 50% variable metabolites across cell lines of distinct cancer types (Figure S2F), followed by enrichment analysis via MetaboAnalyst ^30^, we observed a significant correlation between metabolomics-based variable pathways and gene expression-based variable pathways (Pearson’s *r* = 0.4, *P* = 0.027) (Figure S2G).

Together, our systematic analysis of metabolic gene expression data across many cell lines and tumors as well as metabolite abundance data on cell lines showed that variability in metabolic pathways could be explained at least partly by differences in cancer type or lineage. Linoleic acid, phenylalanine, and ascorbate metabolism were identified among pathways with substantial cancer type-associated heterogeneities, while variability in some other pathways, such as Oxphos and TCA cycle, could not be explained by differences in cancer type.

### Oxphos state exhibits substantial cell line-to-cell line and cell-to-cell heterogeneity independent of cancer type and growth condition

Focusing on each cancer type separately, we then sought to assess the extent of heterogeneity in each metabolic pathway among individual cell lines. To identify heterogeneous metabolic pathways in cell lines within each cancer type, we first performed PCA on the gene expression dataset from cell lines of that cancer type. We computed metabolic gene variability scores as the absolute sum of the PCA loading values across the top PCs accounting for at least 80% of the variance in each dataset. We then applied pre-ranked gene set enrichment analysis (GSEA) to the ranked lists of metabolic gene variability scores. GSEA revealed cell lines of the same cancer type with significant variability in genes found in 25 metabolic pathways (Figure 3A). Hierarchical clustering revealed several cancer types exhibiting more profound cell line-to-cell line heterogeneity in some pathways than others. We thus ranked metabolic pathways according to the number of cancer types showing significant cell line-to-cell line heterogeneity (Figure 3B). The top three pathways with the highest level of cell line-to-cell line heterogeneity were retinol metabolism, cytochrome P450 metabolism, and oxidative phosphorylation (Oxphos). The first two were also among the pathways whose relative activity varied substantially from one cancer type to another (Figures 2A and S2C). Surprisingly, however, Oxphos had shown the least amount of variability among cell lines when compared collectively across distinct cancer types (Figures 2A and 2B). Other metabolic pathways showed different trends; ascorbate and aldarate metabolism, for example, was largely variable across diverse cancer types as well as among individual cell lines of some cancer types (Figures 2A, S2C and 3A). These distinct patterns were also clear when we visualized the distributions of the median and interquartile range (IQR) values for the cumulative abundance of gene transcripts corresponding to each metabolic pathway (Figures 3C and S3). For example, the median and IQR for the abundance of ascorbate and aldarate metabolism transcripts varied largely (up to 7-fold and 21-fold, respectively) among groups of myeloproliferative neoplasm, AML, melanoma, colorectal adenocarcinoma, and pancreatic adenocarcinoma cell lines. In contrast, the median and IQR for the abundance of Oxphos transcripts varied minimally (<1.5-fold and <1.3-fold, respectively) across the same groups of cell lines that showed significant cell line-to-cell line heterogeneity (Figures 3C). By analyzing single-cell RNA-sequencing data ^31^, we evaluated the single-cell distribution of Oxphos state across individual cells within both Oxphos^High^ and Oxphos^Low^ subgroups of cell lines. We observed a high degree of cell-to-cell heterogeneity, spanning the entire spectrum of Oxphos state, in almost all tested cell lines (Figure 3D). What varied between Oxphos^High^ and Oxphos^Low^ cell lines was the frequency of Oxphos^High^ and Oxphos^Low^ cells, rather than a whole-population shift in Oxphos state.

**Figure 3.**
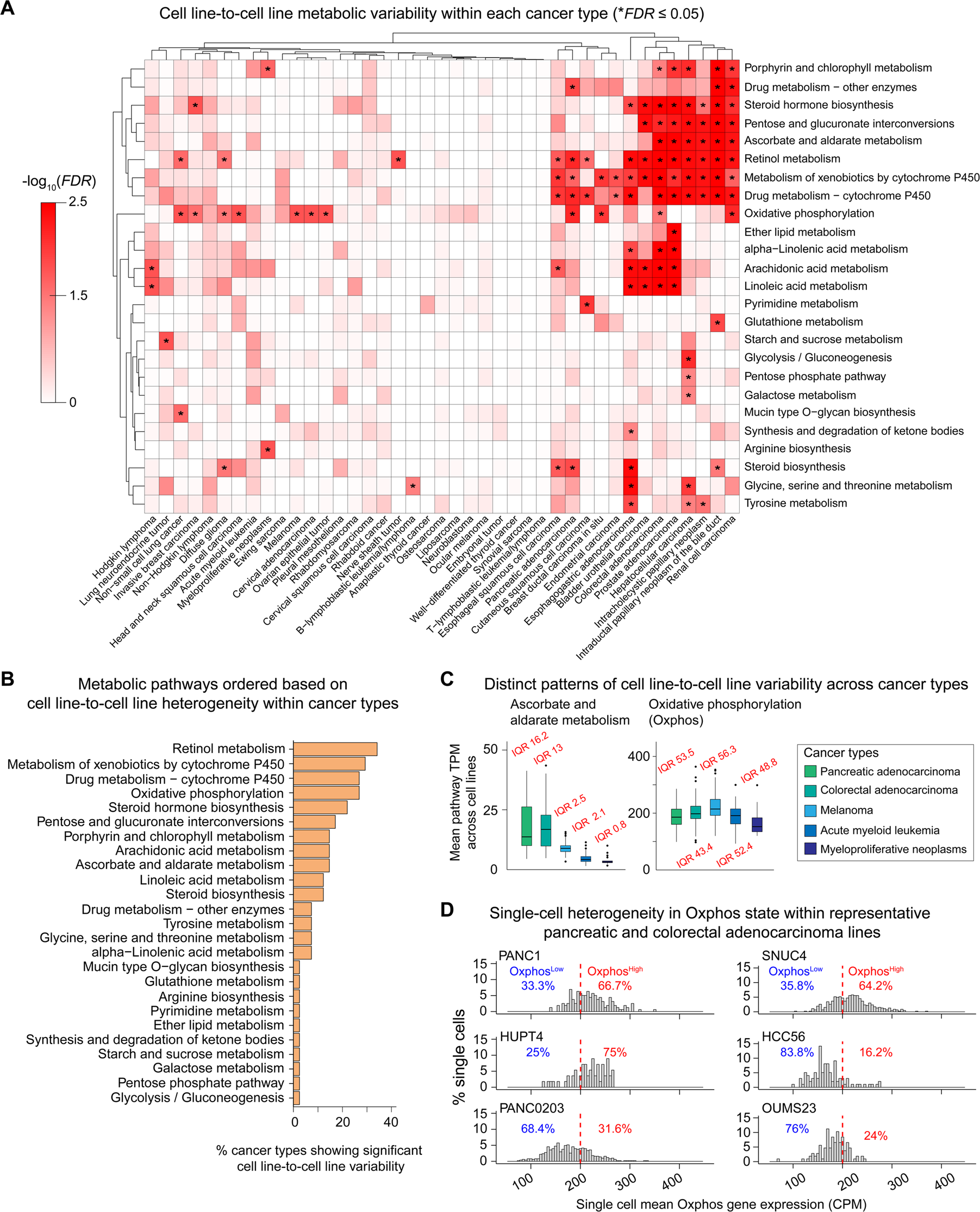
Oxphos state exhibits substantial cell line-to-cell line heterogeneity independent of cancer type. **(A)** Metabolic pathways enriched in genes with highest contribution (determined by PCA) to the metabolic heterogeneities among individual cell lines from different cancer types. Data are clustered based on-log_10_(FDR) values derived from gene set enrichment analysis (GSEA). Pairs of pathway/cancer type with significant cell line-to-cell line heterogeneity (*FDR* ≤ 0.05) are highlighted with an asterisk (*). **(B)** Metabolic pathways ranked according to the percent of cancer types showing significant cell line-to-cell line heterogeneity**. (C)** Distinct patterns of cell line-to-cell line variability in two representative metabolic pathways, ascorbate and aldarate metabolism (left) and oxidative phosphorylation (right), across indicated cancer types. Mean pathway transcript per million (TPM) levels, their median and interquartile ranges (IQR) across cell lines are highlighted. **(D)** Single-cell analysis of Oxphos state based on single-cell RNA sequencing data from representative pancreatic adenocarcinoma (left) and colorectal adenocarcinoma cell lines (right). Mean counts per million (CPM) scores for Oxphos genes were calculated for each single cell. The arbitrary threshold of 200 CPM (red dotted line) was used to separate Oxphos^High^ and Oxphos^Low^ cells.

Cancer cell lines are grown in different culture conditions (e.g., adherent versus suspension) with various growth media that may influence their metabolic states and dependencies ^13,14,32^. We thus quantified the statistical contribution of growth conditions toward the observed cell line-to-cell line heterogeneity in metabolic pathways. We used the same approach as described above to compute relative metabolic pathway activity scores across groups of cell lines and determined the strength of metabolic pathway associations with each of the 21 chemically distinct media compositions (Figures S4A and 4A) or 4 different cell culture conditions (Figures S4B and S4C). Among the pathways whose activity was most affected by growth condition were linoleic acid metabolism, retinol metabolism, glycosaminoglycan biosynthesis, ascorbate and aldarate metabolism (Figures 4B and S4D). On the other hand, variations in a subset of pathways, including Oxphos and TCA cycle exhibited little association with either growth media composition or culture condition (Figures 4B and S4D). We also did not see a significant association between Oxphos state and doubling time of cell lines (Figure S4E). These results suggest a dominant role for cell-intrinsic factors in driving the observed heterogeneity in Oxphos state across cell lines. Cell line-to-cell line variability in pathways such as linoleic acid and retinol metabolism, however, could be attributed at least partly to differences in cellular growth condition. To test the generalizability of this idea, we analyzed metabolic gene expression data for an independent set of cell lines from the Sanger Cell Lines Project ^33^. Most genes showed consistent expression profiles across the 658 cell lines shared between CCLE and Sanger datasets (Figure S4F). However, expression profiles for ∼20% of metabolic genes showed a Pearson’s correlation of <0.5 between the two datasets (Figure S4F). The top pathways represented by these genes were those whose activity we found to be impacted most by growth condition, such as linoleic acid, retinol, and phenylalanine metabolism (Figure S4G). Oxphos, on the other hand, stood out consistently among the most variable pathways across individual cell lines within various cancer types in both Sanger and CCLE datasets (Figures S4H and 3A).

**Figure 4.**
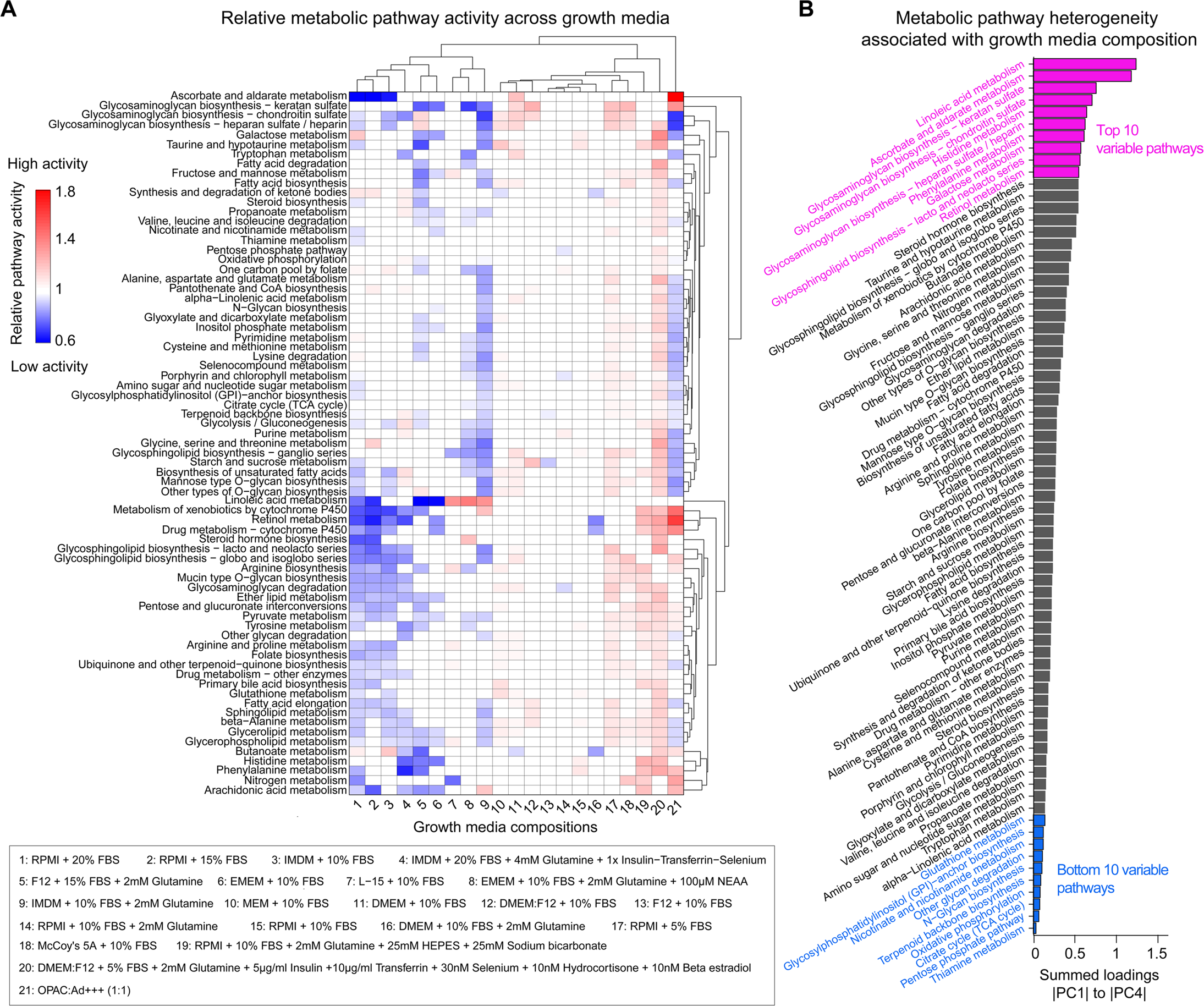
Metabolic pathway analysis of cancer cell lines reveals variations in the activity of pathways associated with growth media. **(A)** Hierarchical clustering of relative activity scores for 73 significantly variable metabolic pathways across 21 chemically distinct growth media. A metabolic pathway activity score < 1 for a growth medium represents reduced pathway activity in that growth medium in comparison with the average pathway activity across all growth media; scores > 1 represent increased activity; and a score of 1 represents activity levels equivalent to the average over all growth media. **(B)** Ranking of metabolic pathways based on the extent to which their heterogeneity is associated with growth media, evaluated by computing the absolute sum of PCA loadings for each pathway over the first four principal components.

### Multivariate modeling uncovers pan-cancer gene vulnerabilities associated with Oxphos state

As shown above, Oxphos was among the most variable pathways across individual cell lines in multiple cancer types and its variability could not be explained solely by differences in growth condition. Therefore, we set out to leverage intrinsic Oxphos heterogeneity to uncover Oxphos state-associated gene vulnerabilities. We hypothesized that novel therapeutic targets specific to Oxphos^High^ and Oxphos^Low^ states could be identified through conditional synthetic lethality. To test this hypothesis, we first analyzed the data from genome-wide DepMap CRISPR knockout screen across 495 CCLE cell lines ^12^, representing 11 cancer types with significant cell line-to-cell line heterogeneity in Oxphos, including non-small cell lung cancer (NSCLC), invasive breast carcinoma (IBC), diffuse glioma (DG), head and neck squamous cell carcinoma (HNSCC), melanoma, cervical adenocarcinoma (CAC), ovarian epithelial tumor (OET), pancreatic adenocarcinoma (PAC), breast ductal carcinoma in situ (BDCIS), colorectal adenocarcinoma (CRAC), and renal cell carcinoma (RCC). We used DepMap gene dependency scores as measures of the effect of 17,202 gene knockouts on viability of each cell line, where a score of 0 indicates no inhibitory effect (corresponding to a non-essential gene), and 1 indicates complete inhibitory effect (corresponding to an essential gene) ^34^. To systematically identify gene vulnerabilities that were most strongly related to Oxphos state, we performed feature selection and multivariate modeling.

We first calculated Oxphos state scores across the cell lines, defined as the average of Z scores for the expression levels of 113 Oxphos genes. To amplify the impact of Oxphos^High^ and Oxphos^Low^ groups of cell lines in feature selection, we ordered cell lines based on their Oxphos state scores and included only the top and bottom 33 percentiles, resulting in 328 cell lines (Figure S5A). As the first step of feature selection, we removed genes with dependency scores that varied minimally across Oxphos^High^ and Oxphos^Low^ cell lines, narrowing down the list of genes to 3,624 (Figure S5B). We then used elastic net regularization to select a subset of the remaining genes based on their association with the Oxphos state score. We optimized the parameters of elastic net model and trained it over 150 iterations (using 10-fold cross validation) using a randomly selected group of 296 cell lines out of 328 cell lines, leaving the other 32 cell lines to be used as a test set for independent validation (Figures S5C and S5D). 200 genes appeared in at least 50% of all elastic net iterations (Figure S5E). We imported these genes into Enrichr to search for potential enrichment of cellular components, biological processes, and pathways associated with the selected gene vulnerabilities (Figures 5A and S5F). We found Oxphos^High^ cell lines were vulnerable to gene knockouts associated with mitochondrial membrane, respiratory electron transport chain (ETC) and histone acetyl transferase complexes. Oxphos^Low^ cell lines, on the other hand, were vulnerable to knockouts associated with Rho GTPase signaling, focal adhesion, cell-matrix junctions, lysosomes, and the trans-Golgi network. An independent but similar analysis on the Sanger dataset ^35^ (including feature selection followed by enrichment analysis) revealed similar patterns of vulnerability in Oxphos^High^ versus Oxphos^Low^ cell lines, although the dataset included ∼50% fewer cell lines (Figures S5G and S5H).

**Figure 5.**
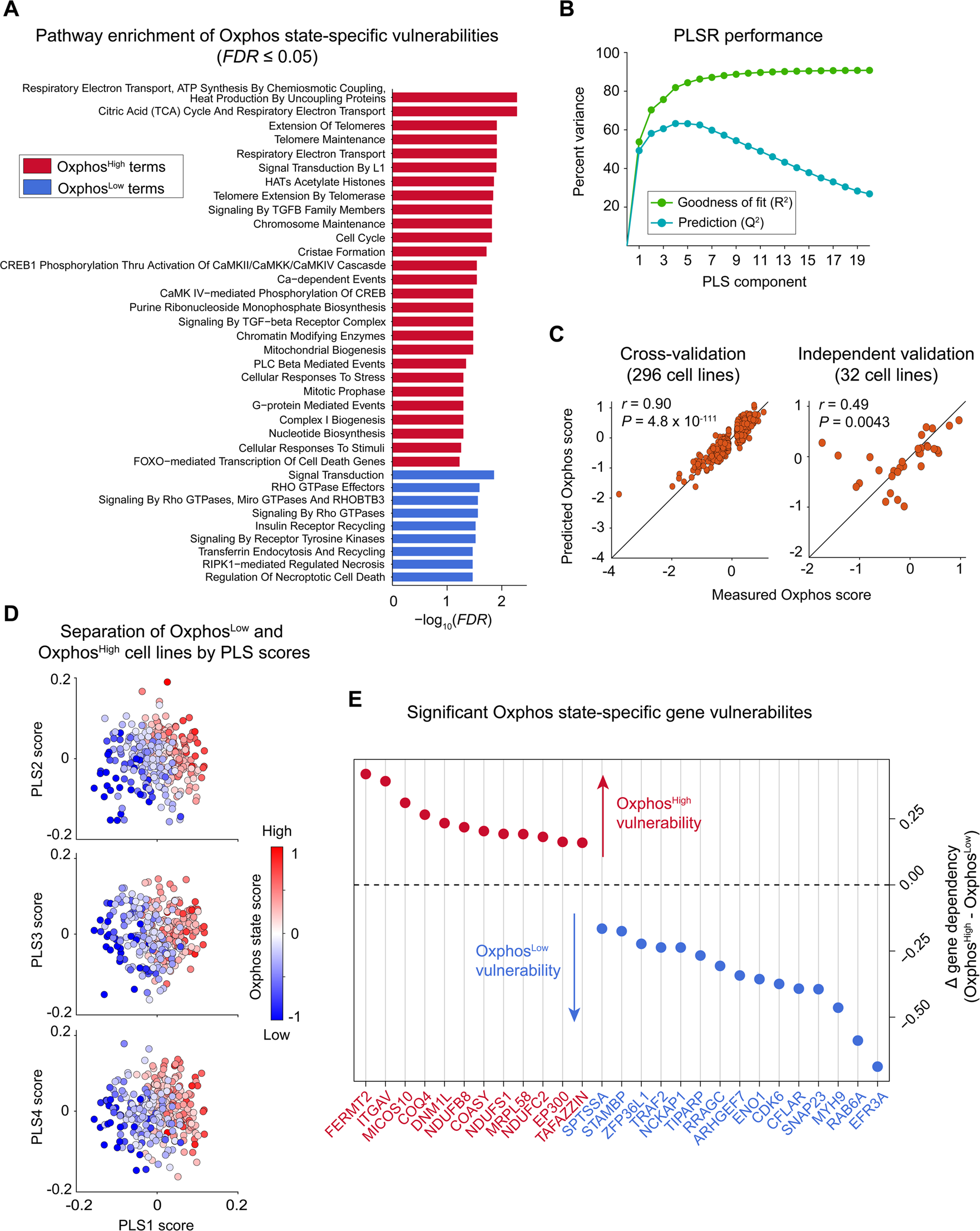
Multivariate modeling uncovers pan-cancer gene vulnerabilities associated with Oxphos state. **(A)** The top enriched biochemical pathways (from Reactome Pathway Database) associated with gene vulnerability features selected by elastic net regularization (*FDR* ≤ 0.05). Genes with positive and negative elastic net coefficients were used to infer vulnerabilities in Oxphos^High^ and Oxphos^Low^ cell lines, respectively. **(B)** Performance of the partial least squares (PLSR) model evaluated by variance in Oxphos state scores explained (R^2^) or predicted based on leave-one-out cross-validation (Q^2^) with increasing number of PLS components. **(C)** Comparison between Oxphos state scores and predicted scores by PLSR for the training set of 296 cell lines (left) and independent test set of 32 cell lines (right). Two-sided Pearson’s correlation analysis was performed between Oxphos state scores and PLSR predicted scores. **(D)** PLSR scores colored according to their Oxphos state score. **(E)** The list of pan-cancer gene vulnerabilities strongly associated with the Oxphos^High^ or Oxphos^Low^ state. For each gene, the median difference between dependency (Δ gene dependency) for Oxphos^High^ and Oxphos^Low^ subgroups of cell lines are shown.

To quantify the relative importance of 200 gene dependencies in predicting the Oxphos state of each cell line, we built a partial least-square regression (PLSR) model ^36,37^. The overall performance of the PLSR model was evaluated based on the percentage of variance in Oxphos state scores explained (R^2^) or predicted (Q^2^) by the variance in gene dependency scores (Figure 5B). The model revealed high performance with R^2^ of 81.9% and Q^2^ of 63.3% (using leave-one-out cross-validation) for the first four PLS components. To further evaluate the model performance, we correlated the true Oxphos state scores and the state scores predicted (based on leave-one-out) by the PLSR model, which led to a Pearson’s correlation coefficient of 0.9 (*P* = 4.8 × 10^-111^) across 296 cell lines (Figure 5C; left panel). To independently validate the model, we used the test set of 32 CCLE cell lines that were not included in either elastic net regularization or PLSR training steps (Figure S5C). The trained PLSR model was able to predict Oxphos state scores in the test set with a Pearson’s correlation coefficient of 0.49 (*P* = 0.004) (Figure 5C; right panel).

The high performance of the PLSR model suggests that variations in the knockout effects of predictive genes can explain variability in Oxphos state across heterogeneous cell lines. This was also evident from the observation that cell lines with high or low Oxphos scores could be efficiently separated based on their PLSR scores along PLS1 to PLS4 (Figure 5D). To identify those gene vulnerabilities that most significantly predicted the Oxphos state, we calculated the variable importance in projection (VIP) scores along the first four PLS components. We identified 64 genes showing VIP scores of greater than one (Figure S5I). We then used a combination of permutation testing and Pearson’s correlation analysis to identify 53 out of 64 genes, which not only showed significant differences in their dependencies between subgroups of Oxphos^High^ and Oxphos^Low^ cell lines (*P* ≤ 0.05), but also retained their correlations with Oxphos state score when evaluated across the complete list of 495 cell lines (*FDR* ≤ 0.05). Among these genes, we focused on those whose median dependencies between subgroups of Oxphos^High^ and Oxphos^Low^ cell lines were larger than 0.15. This led to a final list of 12 gene vulnerabilities strongly associated with the Oxphos^High^ state and 15 gene vulnerabilities strongly associated with the Oxphos^Low^ state (Figure 5E).

Among the most significant predictors of Oxphos^High^ state were gene dependencies associated with mitochondrial membrane and function, including MICOS10 and TAFAZZIN (which play critical roles in maintaining mitochondrial inner membrane structure and function), COQ4, NDUFB8, NDUFS1 and NDUFC2 (mediators of mitochondrial membrane respiratory chain), DNM1L (a mediator of mitochondrial fission), and MRPL58 (a component of the large mitochondrial ribosome required for mitochondrial gene translation). We also found EP300 (a histone acetyltransferase that regulates transcription via chromatin remodeling) and COASY (an enzyme involved in biosynthesis of coenzyme A, a carrier of acetyl and acyl groups) to be additional Oxphos^High^ state-specific dependencies. On the other hand, the list of Oxphos^Low^ state-specific dependencies included ENO1 (a key glycolytic enzyme), TIPARP (a regulator of glycolytic gene expression through HIF-1α ^38^), and genes involved in Rho and Rab GTPase signaling, such as ARHGEF7, NCKAP1 and MYH9 (involved in cytoskeleton regulation), RAB6A, SNAP23 and EFR3A (involved in vesicular trafficking, membrane fusion and Golgi organization).

### Statistical analysis reveals synthetically lethal associations between Oxphos state, driver mutations and tissue context

Next, we asked whether any of the Oxphos state-associated gene vulnerabilities were enriched more significantly in cell lines originating from specific tissue types or those carrying specific driver mutations. To answer this question, we compared median gene dependency between Oxphos^High^ and Oxphos^Low^ subgroups among cell lines grouped based on cancer type or occurrence of the top 25 commonly mutated driver genes ^39^ (Figure S6A). Among these genes, we focused on the top eleven, which carried mutations in at least 15 tested cell lines, including TP53, KRAS, PIK3CA, BRAF, APC, ARID1A, PTEN, NF1, FAT1, KMT2D, CREBBP (Figure S6B). To assess the statistical significance of the observed differences in gene dependency, we performed permutation testing by randomly shuffling the Oxphos state labels (High and Low) across cell lines participating in each comparison and computing permutation *P* values. We then used *P* ≤ 0.01 as the significance cutoff to identify mutations and tissue types associated with enhanced Oxphos state-specific gene dependencies relative to pan-cancer analysis.

Focusing on Oxphos^High^ state-specific dependencies, we found that mutations in the tumor suppressor PTEN were consistently associated with significant increases in dependency on genes involved in mitochondrial membrane respiratory chain, including COQ4, NDUFB8, NDUFS1 and NDUFC2 (Figures 6A and 6B). KRAS and APC mutations were associated with enhanced dependency on DNM1L (Figure S6C), and the effect of EP300 gene knockout in Oxphos^High^ cell lines was significantly greater in the presence of mutations in chromatin regulators KMT2D or ARID1A (Figure 6C). We also identified cases, where an Oxphos^High^ state-specific gene dependency was enriched in cell lines from a specific cancer type, such as EP300 dependency in melanoma and ovarian epithelial tumor cell lines and MRPL58 dependency in invasive breast carcinoma (IBC) and head and neck squamous cell carcinoma (HNSCC) cell lines (Figures 6A and 6C). Focusing on Oxphos^Low^ state-specific dependencies, we found that multiple gene dependencies associated with cytoskeleton regulation and vesicle trafficking (such as ARHGEF7, RAB6A, NCKAP1) were significantly enriched in pancreatic adenocarcinoma cell lines (Figure 6D). The most frequent oncogenic events correlated with Oxphos^Low^ state-specific gene dependencies were FAT1 mutations that were associated with enhanced dependency on ARHGEF7, ENO1 and ZFP36L1, and ARID1A mutations that were associated with increased dependency on RAB6A, RRAGC, TIPARP and ZFP36L1 (Figures 6D and S6D).

**Figure 6.**
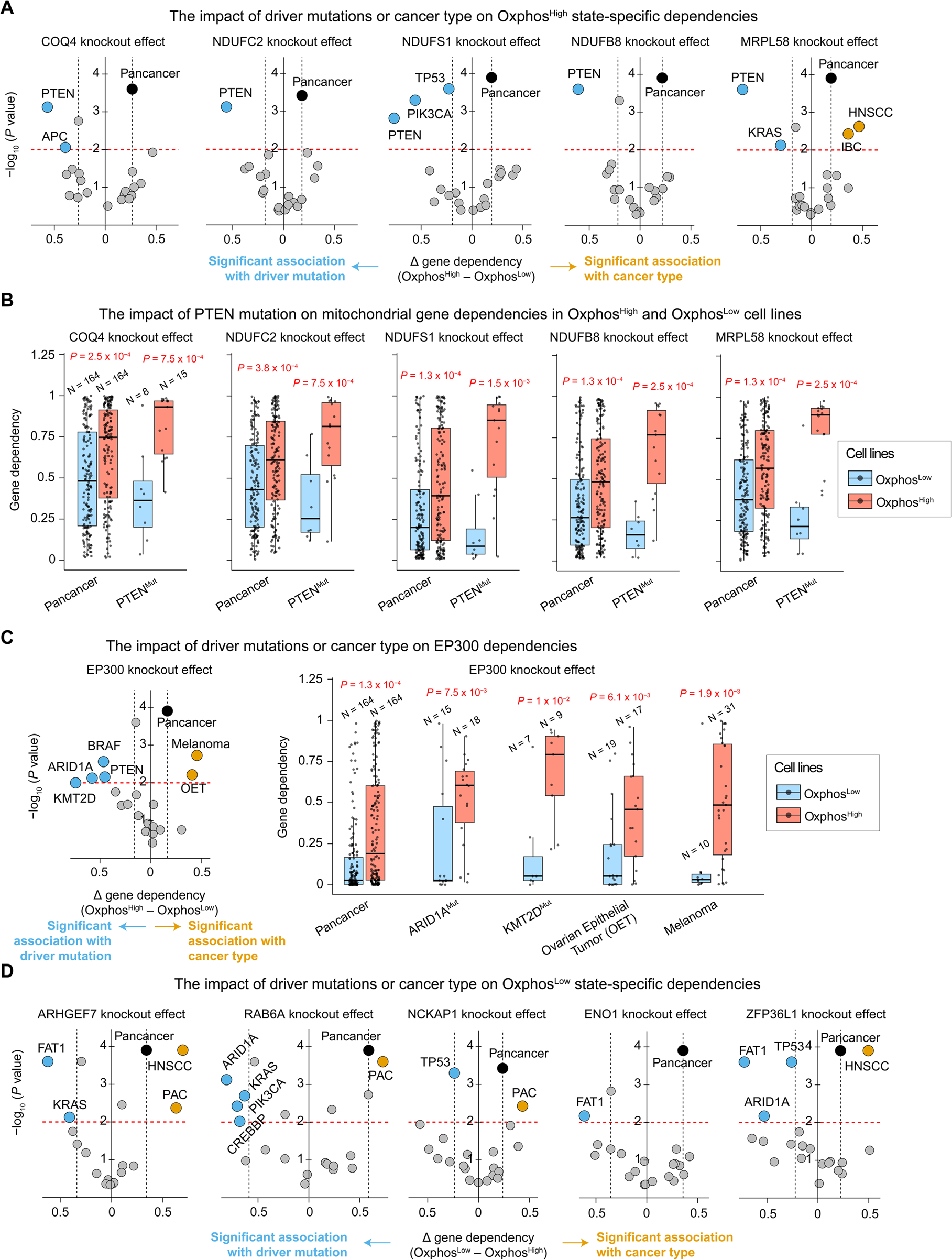
Statistical analysis reveals synthetically lethal associations between Oxphos state, driver mutations and tissue context. **(A)** Statistical enrichment of Oxphos^High^-associated mitochondrial gene vulnerabilities in cell lines associated with specific tissue types or driver mutations. Enrichment is considered significant if the median gene dependency difference (Δ gene dependency) between Oxphos^High^ and Oxphos^Low^ subgroups of cell lines associated with a cancer type or driver mutation is significantly larger than the median gene dependency difference between Oxphos^High^ and Oxphos^Low^ groups regardless of cancer type and mutation (i.e., pan-cancer median difference level highlighted by the black circle and dotted line). Statistical significance of median differences was determined by empirical *P* values computed based on permutation testing (*P* ≤ 0.01; red dotted line). **(B)** The impact of PTEN mutation on mitochondrial gene dependencies in Oxphos^High^ and Oxphos^Low^ cell lines. Statistical significance of median differences was determined by empirical *P* values computed based on permutation testing. **(C)** Statistical enrichment of Oxphos^High^-associated EP300 dependency in cell lines associated with specific tissue types or driver mutations. Statistical significance of median differences was determined by empirical *P* values computed based on permutation testing. **(D)** Statistical enrichment of Oxphos^Low^-associated gene vulnerabilities in cell lines associated with specific tissue types or driver mutations. Statistical significance of median differences was determined by empirical *P* values computed based on permutation testing.

Together, these results suggest that synthetically lethal relationships exist between the tumor metabolic state, driver mutations and specific genes or pathways that could be potentially exploited to selectively target cancer state-specific vulnerabilities. To further explore this idea, we focused our attention on PTEN mutations based on their profound association with Oxphos^High^ state-specific dependencies on mitochondrial membrane structure and electron transport chain (ETC).

### Loss of PTEN predicts increased dependency on mitochondrial respiratory chain in Oxphos^High^ tumor cells

To independently test the impact of damaging mutations in PTEN on Oxphos state-specific dependencies, we used the Cancer Therapeutics Response Portal (CTRP) data to analyze the sensitivity of 799 genetically characterized cancer cell lines to 545 small-molecule probes and drugs ^40^. We used transcriptomics data to define groups of Oxphos^Low^ and Oxphos^High^ cell lines based on whether their Oxphos state scores were ranked within the top or bottom 33 percentiles. We then compared the median sensitivity of Oxphos^Low^ and Oxphos^High^ cell lines to each tested small molecule based on the area of the dose-response curve (AUC) measurements. Within the Oxphos^High^ group, we also compared the median sensitivity of PTEN-wildtype (PTEN^WT^) and PTEN-mutated (PTEN^Mut^) subgroups. Among all small molecules tested, we found oligomycin A, a well-known inhibitor of mitochondrial ATP synthase (complex V) ^41^, to exhibit the strongest differential effect on Oxphos^High^ cell lines relative to Oxphos^Low^ cell lines (Median difference = 0.9; *P* = 0.04), while being more effective in the PTEN^Mut^ subgroup than in PTEN^WT^ cells (Median difference = 1.7; *P* = 0.07) (Figure 7A). Although not statistically significant, neopeltolide, a natural product which was recently discovered as an inhibitor of mitochondrial ATP synthesis ^42^, was also among the top 4 compounds with increased median efficacy in Oxphos^High^ cell lines compared with Oxphos^Low^ cell lines (Median difference = 0.9), while also being more effective in the PTEN^Mut^ subgroup than in PTEN^WT^ cells (Median difference = 0.9). These results are consistent with the evidence from the CRISPR knockout screen data, showing enhanced dependency of PTEN^Mut^/Oxphos^High^ tumor lines on mitochondrial ETC.

**Figure 7.**
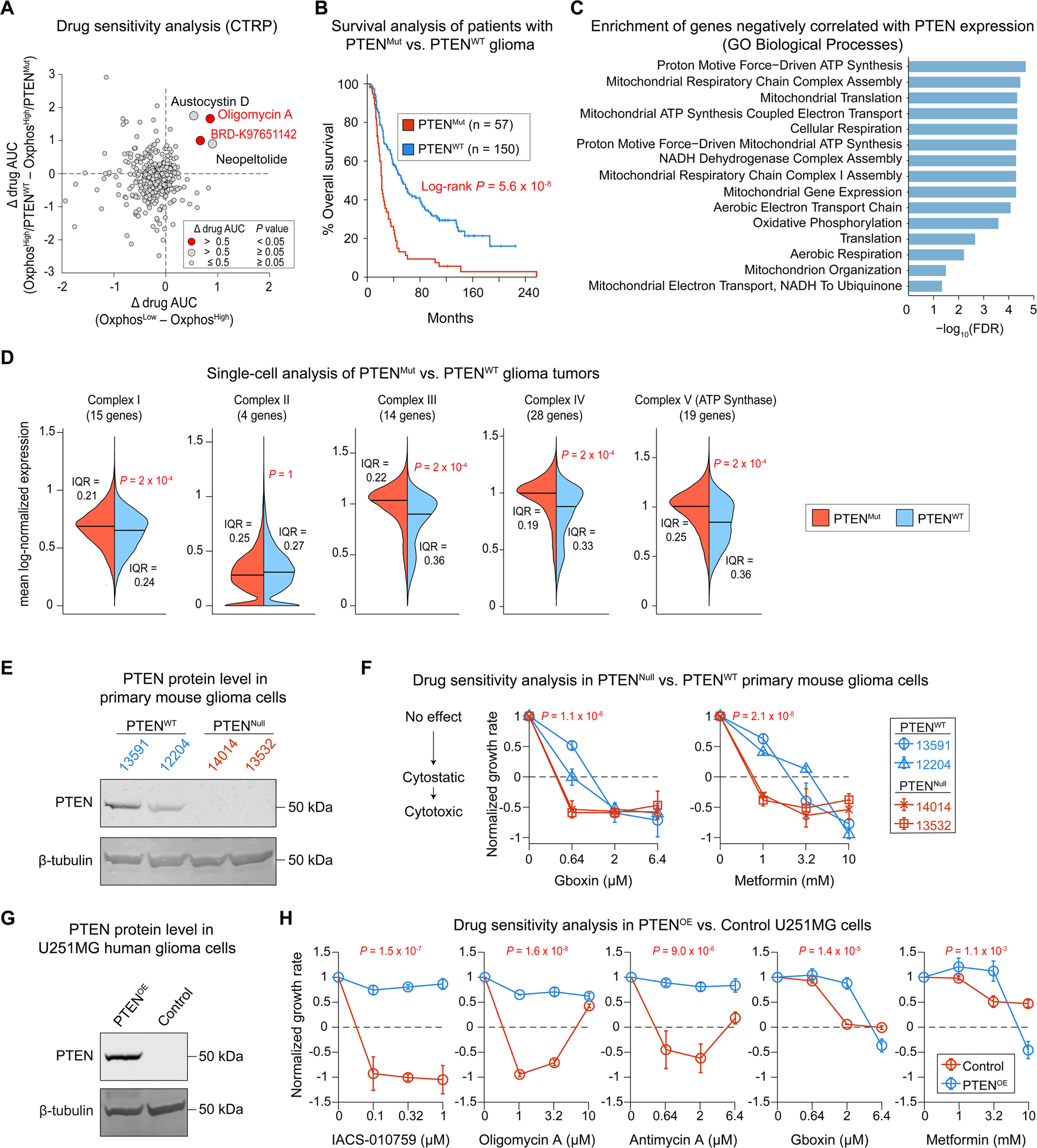
Loss of PTEN predicts increased dependency on mitochondrial respiratory chain in Oxphos^High^ glioma tumor cells. **(A)** Analysis of the Cancer Therapeutics Response Portal (CTRP) data, including sensitivity measurements (evaluated based on area under the dose response curve; AUC) for 545 small-molecule probes and drugs in 799 cell lines. For each compound, the difference in median sensitivity (Δ drug AUC) between Oxphos^Low^ and Oxphos^High^ subgroups of cell lines is shown against the difference in median sensitivity between Oxphos^High^/PTEN^WT^ and Oxphos^High^/PTEN^Mut^ subgroups. Statistical significance was determined using one-sided Wilcoxon rank sum test. **(B)** Overall survival analysis of glioma patients with PTEN-mutated (PTEN^Mut^) or PTEN-wildtype (PTEN^WT^) tumors. Statistical significance was determined by two-sided log-rank (Mantel-Cox) test. **(C)** The top Gene Ontology (GO) biological processes (*FDR* ≤ 0.05) associated with genes whose expression negatively correlated with PTEN mRNA levels across 79 glioma tumors. **(D)** Single-cell analysis of PTEN^Mut^ versus PTEN^WT^ glioma tumors. Mean log-normalized levels of transcripts representing individual components of the electron transport chain (ETC), including Complex I-IV and ATP synthase, between PTEN^Mut^ (n = 10,268) and PTEN^WT^ (n = 23,229) malignant cells. Statistical significance was determined by one-sided, permutation test. **(E)** PTEN protein levels assessed by Western blotting in two PTEN^Null^ (14014 and 13532) and two PTEN^WT^ (13591 and 12204) primary mouse glioma cell lines. **(F)** Dose-dependent changes in normalized growth rates of PTEN^Null^ and PTEN^WT^ primary mouse glioma cell lines following treatment with Gboxin and metformin. The average net growth rates were calculated from measurements of live cell count at three timepoints (including 0, 96, and 168 hours). Statistical significance of the effect of each drug on PTEN^Null^ vs. PTEN^WT^ cell lines was determined by two-way ANOVA, conducted specifically on the second lowest dose of each drug (i.e., 0.64 µM for Gboxin and 1 mM for metformin). **(G)** PTEN protein levels assessed by Western blotting in stable U251MG human glioma cells overexpressing PTEN (PTEN^OE^) or Vector control (Control). **(H)** Normalized growth rates measured in U251MG cells overexpressing PTEN (PTEN^OE^) or Vector control (Control) following treatment with indicated concentrations of IACS-010759, oligomycin A, antimycin A, Gboxin, and metformin. The average net growth rates were calculated from measurements of live cell count at three timepoints (including 0, 96, and 144 hours). Statistical significance of the effect of each drug on PTEN^OE^ vs. Control cells was determined by two-way ANOVA, conducted specifically on the middle two doses of each drug (i.e., 0.1 and 0.32 µM for IACS-010759; 1 and 3.2 µM for oligomycin A; 0.64 and 2 µM for antimycin A and Gboxin; 1 and 3.2 mM for metformin). Drug sensitivity data in **F** and **H** are presented as mean values ± SD across 3 replicates.

To further explore the role of PTEN loss in tumor cells’ dependency on mitochondrial ETC, we turned our focus on diffuse gliomas, the most common malignant adult brain cancer, in which genetic loss of PTEN expression is frequently observed ^43^. Multiple studies have reported the sensitivity of glioma cell lines and tumors to Oxphos inhibitors that block mitochondrial ETC protein complexes *in vitro* and *in vivo* ^44–46^. However, little is known about how the genetic background of glioma tumors might influence such metabolic dependency. To explore this, we analyzed DNA-and RNA-sequencing data and matched clinical annotation for glioma patients across glioma subtypes assembled by the GLASS consortium ^47^. Tumors from 64 out of 222 patients (i.e., ∼29% of patients) carried PTEN mutations, including truncating, missense, in-frame, and splice-site mutations. The presence of such mutations in tumors were associated with significantly shorter overall survival in glioma patients (Figure 7B). To infer how PTEN expression might be associated with glioma tumor state, we performed transcriptome-wide co-expression analysis across 79 tumor samples for which RNA-sequencing data were available. We used Spearman’s correlation analysis to rank transcripts based on the association of their abundance with PTEN expression (Figure S7A). We then imported the list of 571 transcripts that significantly and negatively correlated with PTEN expression (Spearman’s *r* < 0; *FDR* ≤ 0.05) into Enrichr to identify cellular components and biological processes associated uniquely with loss of PTEN. The most significantly enriched components and biological processes were mitochondrial membrane, mitochondrial respiratory chain complex, proton-transporting ATP synthase complex and cellular respiration (Figures 7C, S7B, S7C and S7D). These results suggest that glioma tumors with loss of PTEN expression exhibit a higher expression of components of the mitochondrial ETC and ATP synthase complex.

To further test the association of PTEN loss in malignant cells with mitochondrial ETC expression in the same cells, we analyzed single-cell RNA-sequencing data of genetically profiled patient-derived glioblastoma tumors, including 10,268 PTEN^Mut^ tumor cells isolated from a group of 6 patients and 23,229 PTEN^WT^ tumor cells from a group of 20 patients ^48^. We then compared the abundance of transcripts representing individual components of the ETC, including Complex I-IV and ATP synthase (Complex V), between PTEN^Mut^ and PTEN^WT^ cells. We observed significantly higher expression levels for Complex I, III, IV and ATP synthase genes among PTEN^Mut^ cells and a lower level of cell-to-cell variability in their expression (as evaluated by IQR) in comparison with PTEN^WT^ cells (Figure 7D). These results reveal a notable upward shift in the expression of mitochondrial respiratory chain complex genes among PTEN^Mut^ glioma cells, that is consistent with their enhanced dependency on ETC in comparison with PTEN^WT^ tumor cells.

To complement our computational data analysis with experimental validation, we compared the sensitivity of two PTEN^WT^ and two PTEN^Null^ primary mouse glioma cell lines (Figure 7E) to metformin and Gboxin, two ETC inhibitors that were previously shown to be effective against glioma tumors ^44,46^. All cell lines were cultured in serum-free conditions to maintain their biological properties ^49^. Both metformin (an inhibitor of complex I) and Gboxin (an inhibitor of mitochondrial ATP synthase) were able to reduce cellular growth and induce tumor cell killing in a dose-dependent manner in all four cell lines tested (Figure S7E). However, PTEN^Null^ cell lines (14014 and 13532) showed greater sensitivity to significantly smaller doses of both compounds in comparison with PTEN^WT^ cell lines (13591 and 12204) (Figure 7F). To directly validate the connection between PTEN loss and dependency on mitochondrial respiratory chain in an isogenic context, we also evaluated the sensitivity of a PTEN^Null^ human glioblastoma cell line, U251MG, to ETC inhibitors following PTEN rescue via lentiviral overexpression (Figure 7G). In comparison with PTEN^Null^ cells expressing control vector (Control), PTEN-overexpressing (PTEN^OE^) cells exhibited significantly less potent responses to not only Gboxin and metformin, but also IACS-010759 (a clinical-grade inhibitor of complex I) ^45^, oligomycin A (an inhibitor of mitochondrial ATP synthase), and antimycin A (and inhibitor of complex III) (Figures S7F-S7H and 7H). Overall, these data agree with our model predictions and follow-up analysis, confirming a key role for PTEN loss in mediating the increased dependency of glioma tumor cells on mitochondrial respiratory chain.

## Discussion

In this paper, we used a combination of unsupervised and supervised multivariate analysis approaches to integrate transcriptomics data from hundreds of cancer cell lines with their gene dependency scores and mutation profiles, and thereby predict metabolic state-specific vulnerabilities across various tumor lineages and common oncogenic contexts. Our systematic analysis complements previous studies ^11,13,15,16^ by revealing the extent to which heterogeneity in diverse metabolic pathways across cancer cell lines can be explained by differences in their growth environment, their tissue of origin and developmental lineage, or potentially other cell-intrinsic mechanisms. Furthermore, we uncover synthetically lethal associations between the cancer cells’ metabolic state, their driver mutations, and potentially actionable biological targets, that could guide future mechanistic studies toward more precise and context-specific therapeutic opportunities.

We used transcriptomics data to infer the activity of metabolic pathways across more than a thousand cell lines. Our analysis captured key variations in inferred metabolic pathway activities that were consistent with those derived from metabolomics studies of cell lines grown in identical conditions. For example, our analysis highlighted previously reported metabolic differences between hematopoietic and non-hematopoietic cancers as well as other metabolic variations associated with cancer types ^11,15,16^. We also unmasked metabolic similarities among cell lines derived from cancers with related development lineage but in distinct anatomic positions. For example, cell lines of the lung neuroendocrine lineage (represented mostly by the small cell lung cancer cell lines) metabolically clustered more closely with neuroblastoma cell lines rather than with non-small cell lung cancer lines; esophageal squamous cell carcinoma lines also clustered with head and neck squamous cell carcinoma lines rather than with esophagogastric adenocarcinoma lines; and melanoma cell lines of cutaneous and ocular type clustered together. These examples are consistent with previous reports indicating strong associations between metabolic programs and epigenetically regulated tissue lineage and differentiation programs ^15,50,51^, and further highlight the need for integrative multi-omics approaches to build a comprehensive picture of tumor metabolism ^20^.

Consistent with data from the metabolomics studies ^15^, we also observed that pathways associated with lipid, amino acid and carbohydrate metabolism as well as mitochondrial pathways such as Oxphos accounted for the majority of metabolic heterogeneity among cancer cell lines. For many of these pathways, the observed heterogeneity was highly associated with either cancer type and/or growth media. However, cell line-to-cell line heterogeneity in Oxphos appeared to be mostly independent of cancer type. Focusing on cancer cell lines within each cancer type separately, we found Oxphos was highly variable across individual cell lines in at least 11 major cancer types with little association with growth media composition. We found that the Oxphos state in these cell lines was associated with significant and reproducible gene dependencies. Major Oxphos^Low^ state-specific dependencies included genes involved in cytoskeleton regulation, vesicular trafficking, Golgi organization and membrane fusion. These results suggest that alterations in cellular energy status could influence tumor cell dependency on the function of other subcellular organelles such as the Golgi apparatus, and support the idea to target such organelles as potentially selective vulnerabilities associated with tumor metabolic state ^19^. On the other hand, Oxphos^High^ state-specific vulnerabilities included genes encoding enzymes that mediate the mitochondrial membrane respiratory chain and ATP synthesis, as well as genes with key roles in the control and maintenance of mitochondrial shape, crista junctions, and architecture. These findings are consistent with various evidences, indicating a tight connection between mitochondrial network structure and bioenergetics ^22,52,53^, and suggest that mitochondrial network dynamics are linked to metabolic state-specific therapeutic vulnerabilities ^54^.

Finally, our analysis revealed that Oxphos state-specific gene dependencies were enhanced in the presence of some common driver mutations or in specific tissue lineages. We found, for example, that loss of tumor suppressor PTEN was associated with the increased dependency of Oxphos^High^ tumor cells on mitochondrial respiratory chain and their enhanced sensitivity to pharmacological inhibitors of ETC. This finding is also supported by a previous report pointing to increased sensitivity of PTEN-deficient fibroblasts to inhibition of mitochondrial complex I in comparison with PTEN-wildtype cells ^55^. These observations may be especially significant given the narrow therapeutic index of recently developed complex I inhibitors such as IACS-010759 in clinical trials ^56,57^, and highlight the importance of identifying specific subsets of patients that may benefit from lower doses of such therapies. Importantly, loss of PTEN mutations are prevalent and predict poor survival in multiple cancers, including gliomas ^47^. Our transcriptomics analysis of patient-derived glioma tumors revealed a strong association between loss of PTEN and increased expression of genes encoding mitochondrial ETC. These findings are consistent with a recent discovery of a glioma subtype signified by a substantial increase in Oxphos and mitochondrial function and enhanced sensitivity to inhibitors of mitochondrial protein translation and ETC ^58^. Multiple reports have demonstrated the increased efficacy of these inhibitors against high-grade glioma tumors both *in vitro* and *in vivo* ^44–46,59^, providing a promising path toward their clinical evaluation individually or in combination therapies ^60^. In particular, an ongoing phase II trial is testing the efficacy of mitochondrial complex I inhibitor metformin in combination with temozolomide and radiotherapy in patients with tumors that express an Oxphos^High^ signature at diagnosis (NCT04945148). Findings from our study may inform this trial and other clinical studies of mitochondrial inhibitors (e.g., NCT02780024, NCT05929495, NCT05183204, NCT05824559) by proposing that PTEN loss, in addition to the tumor Oxphos state, may serve as a biomarker of therapeutic response. Future studies may also leverage these findings to elucidate the mechanistic basis of the identified synthetically lethal associations and further evaluate their clinical relevance and therapeutic potential in the presence of key microenvironmental players.

### Limitations of the study

One notable limitation of this study is that it primarily relies on transcriptomics data and gene set enrichment analysis to infer the activity of metabolic pathways. Although the large-scale availability of such data at bulk and single-cell levels across thousands of cell lines and tumors provides an enormous opportunity for systematic studies, they do not directly account for metabolic changes at the flux level or abundance of metabolites. Future multi-variate studies, leveraging integrated multi-omic measurements (including metabolomics) in the same tumor cells ^61–63^, will provide additional insights into the regulation of metabolic heterogeneities in tumors and their variation with genetic, tissue, and microenvironmental context. Another limitation is that our analysis of metabolic state-specific vulnerabilities for each cancer type is influenced by the number of CCLE cell lines from that cancer type with available transcriptomics and gene dependency data. Therefore, cell lines of certain cancer types (such as prostate adenocarcinoma) are underrepresented in our analysis, for different reasons such as biological relevance of such cell lines or technical challenges associated with their culture.

## STAR⍰Methods

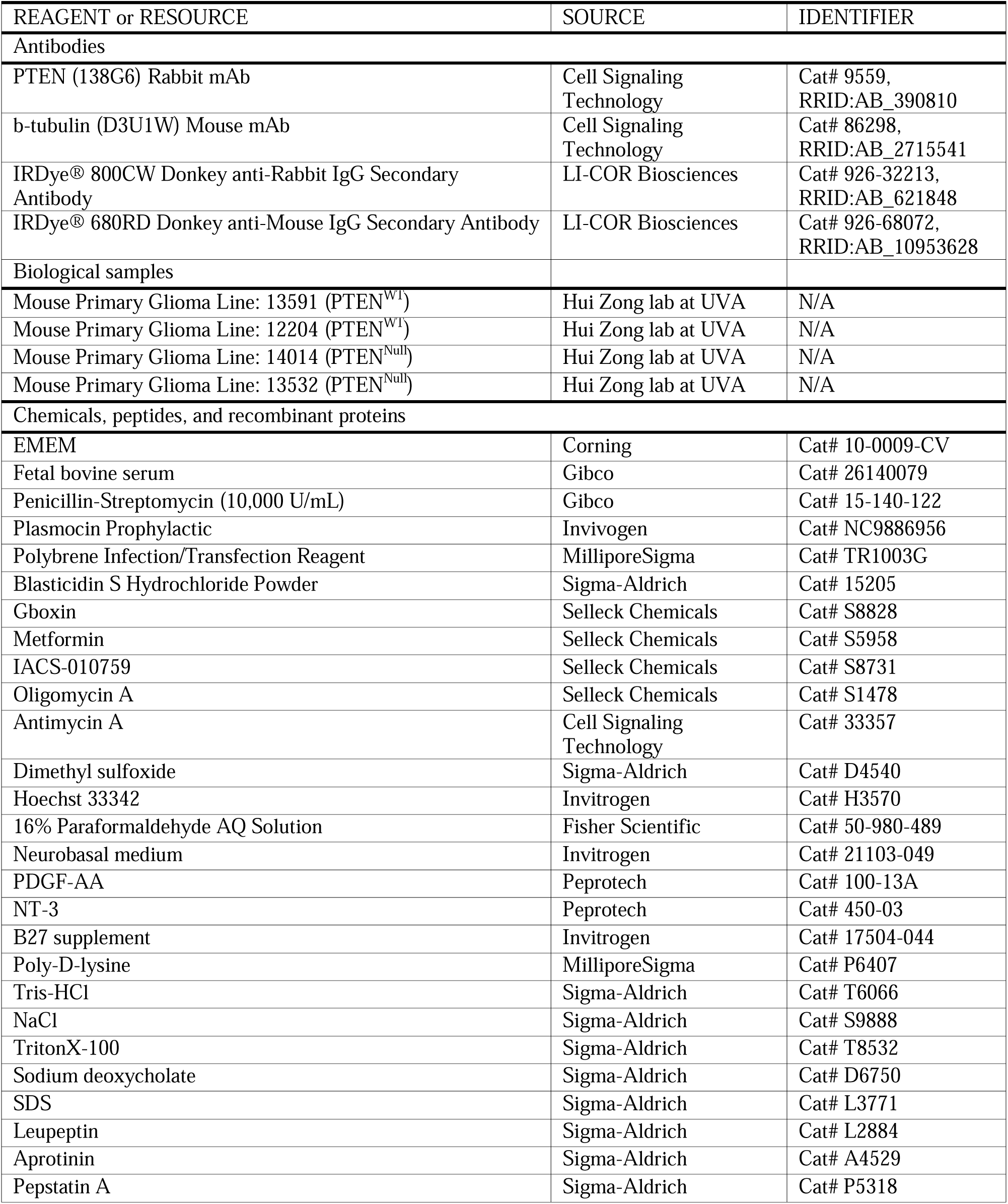
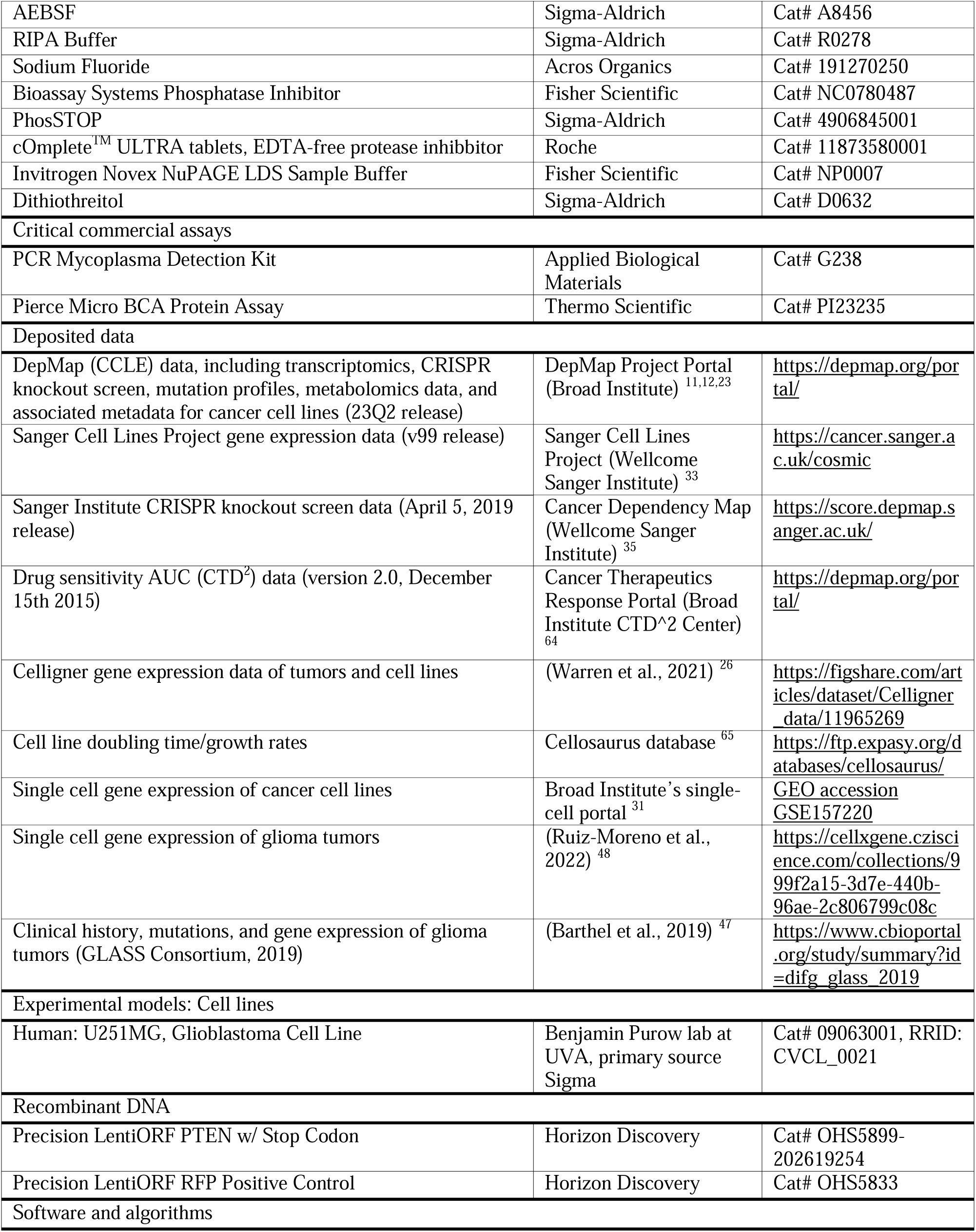
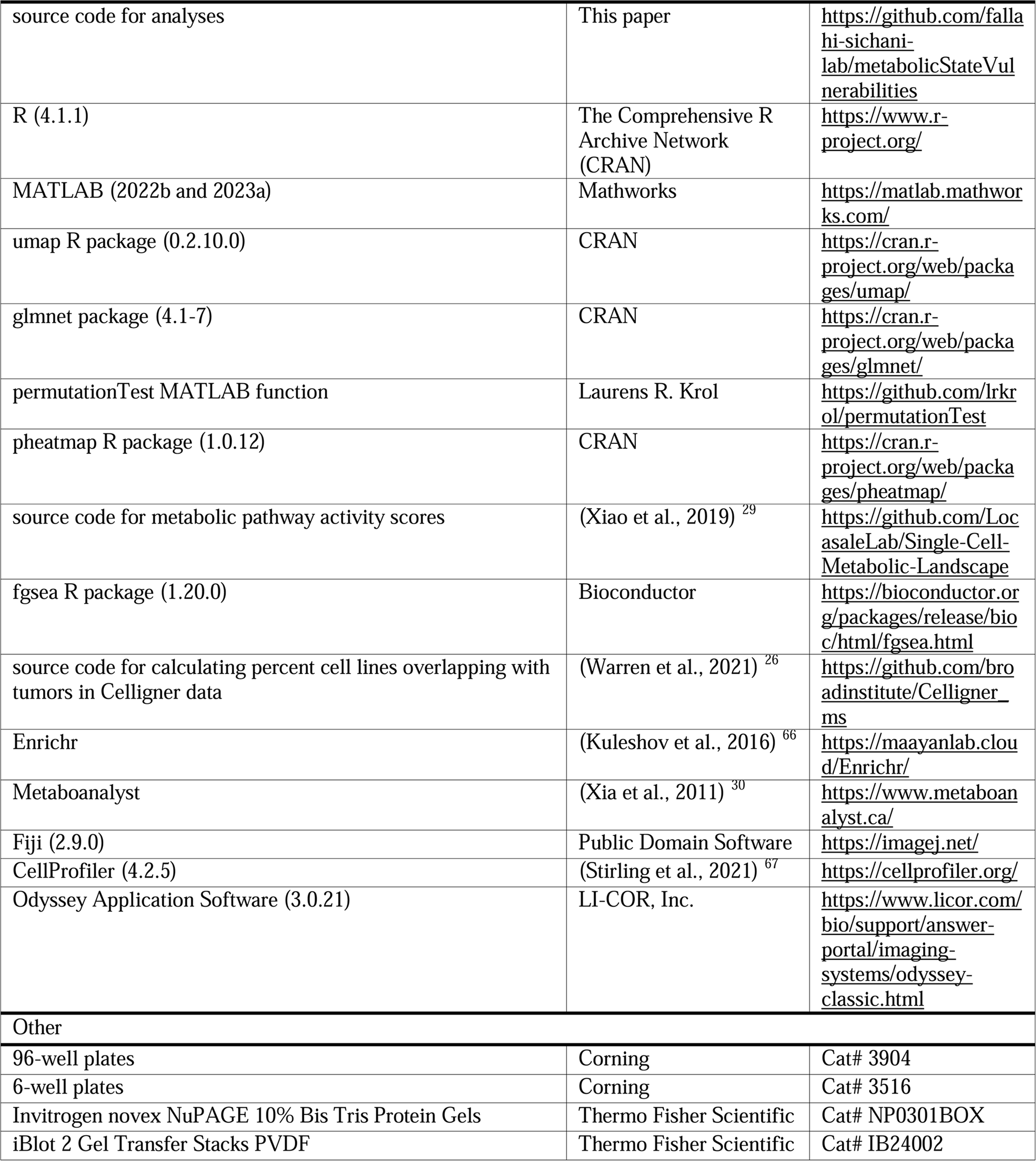
Key resources table.

## Resource availability

### Lead contact

Further information and request for resources should be directed to and will be filled by the lead contact, Mohammad Fallahi-Sichani.

### Materials availability

This study did not generate new unique reagents.

### Data and code availability

- Raw experimental data reported in this paper will be shared by the lead contact upon request. This paper analyzes existing, publicly available data. The accession numbers and identifiers for the datasets are listed in the key resources table.
- The original codes for data analysis performed in this paper are publicly available at GitHub: https://github.com/fallahi-sichani-lab/metabolicStateVulnerabilities
- Any additional information required to reanalyze the data reported in this paper is available from the lead contact upon request.

## Experimental model and study participant details

### Human cell lines

The human malignant glioblastoma cell line used in this study, U251MG, was subjected to confirmation by short tandem repeat profiling by ATCC and Mycoplasma testing by PCR Mycoplasma Detection Kit. U251MG cells were grown at 37°C with 5% CO_2_ within a humidified chamber in Eagle’s Minimum Essential Medium supplemented with 10% FBS, 1% penicillin/streptomycin, and 1:5000 Plasmocin.

### Primary mouse glioma cell culture

Primary mouse PTEN^WT^ glioma cells (13591 and 12204) used in this study were derived from previously established glioma mouse models with inactivating mutations in two tumor suppressor genes Trp53 and Nf1 ^68^. PTEN^Null^ cells (14014 and 13532) were derived from the same model with additional PTEN^flox^/PTEN^flox^ alleles ^69^. After collection from each model, glioma tumor mass was diced into 1 mm^3^ pieces and dissociated with salt balance buffer containing Papain at 20 unit/mL. The tumor tissue was incubated in Papain solution at 37°C 5% CO_2_ for 90 min (gently agitated every 15 min). The Papain digestion was stopped with Trypsin inhibitor and tumor pieces were triturated into cell suspension and passed through a 70 mm cell strainer before centrifugation. Cell pellets were suspended in Neurobasal media supplemented with 2% B27, 10 ng/mL PDGF-AA and 1 ng/mL NT-3. Primary glioma cells were maintained on Poly-D-lysine-treated flasks in Neurobasal media supplemented with 2% B27, 10 ng/mL PDGF-AA and 1 ng/mL NT-3. Cells were grown at 37°C with 5% CO_2_ in a humidified chamber.

## Method details

### Transcriptomic analysis and visualization of metabolic heterogeneity in cancer cell lines

To systematically evaluate metabolic similarities and differences across cell lines representing diverse cancer types, we used RNA sequencing data from the Cancer Cell Line Encyclopedia (23Q2 release). Cancer types were selected if 5 or more cell lines were RNA-sequenced, resulting in 1,341 cell lines spanning across 41 cancer types. Log_2_-transformed gene expression data (i.e., log_2_(TPM+1)) was z-scored for all unique KEGG metabolic genes (1,620 genes in 85 metabolic pathways ^29^) across all cell lines. Uniform manifold approximation and projection (UMAP) clustering was performed in R using the *umap* package (0.2.10.0) with z-scored data using the following parameters: nearest neighbor (n_neighbors) = 50, minimum distance (min_dist) = 0.5, and distance metric (metric) = Pearson. To analyze metabolic heterogeneity within hematopoietic cancer cell lines, we performed principal component analysis (PCA) on the 1,620 metabolic genes using the *prcomp* built-in R function across 218 cell lines of myeloid or lymphoid lineages.

### Calculation of pathway activity scores across different cancer types, growth media, cell culture condition

We computed a score representing the activity of each of the 85 metabolic pathways for each of the cancer types (or growth media or cell culture condition) relative to all cancer types (or all growth media or all culture conditions) using a three-step procedure adapted from Xiao *et al* ^29^. In the first step, we calculated the mean expression level (*E_i,j_*) for each of the 1,620 metabolic genes (i.e., the *i*-th metabolic gene) across cell lines within each of the 41 cancer types (i.e., the *j*-th cancer type) or each of the 21 distinct growth medium compositions (i.e., the *j*-th growth medium) or each of the 4 distinct cell culture conditions (i.e., the *j*-th culture condition):

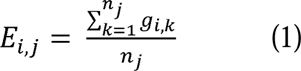

where *n_j_* is the number of cell lines associated with the *j*-th cancer type (or the *j*-th growth medium or *j*-th culture condition), and *g_i,k_* is the log_2_(TPM+1) expression level of the *i*-th gene in the *k*-th cell line in this cancer type (or medium type or culture condition). In the second step, the relative gene expression level of the *i*-th gene in the *j*-th cancer type (or the *j*-th medium type or the *j*-th culture condition) or *r_i,j_* was calculated as the ratio of *E_i,j_*to its average over all cancer types (or all medium types or all culture conditions):

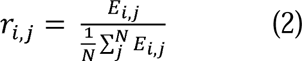

where *N* is the number of cancer types (i.e., 41), medium types (i.e., 21), or culture conditions (i.e., 4). *r_i,j_* values above 1 indicate that the expression level of gene *i* is higher in cancer type *j* (or medium type *j* or culture condition *j*) compared to the average expression level over all cancer types (or all medium types or all culture conditions). In the third step, pathway activity score (*p_t,j_*) for the *t*-th pathway and the *j*-th cancer type (or the *j*-th medium type or the *j*-th culture condition) was calculated as the weighted average of *r_i,j_*over all genes included in the pathway:

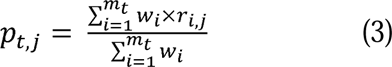

where *m_t_* is the number of genes in pathway *t*, and *w_i_* is a weight factor equal to the reciprocal of the number of pathways that include the *i*-th gene. To avoid pathway activity scores being affected by genes with low and high expression levels, we removed outliers in each pathway defined by genes with expression levels < 0.001 and genes with relative expression levels greater than 3 × 75^th^ percentile or below 1/3 × 25^th^ percentile. We calculated activity scores for pathways that had at least 5 genes after filtering. Statistical significance of lower or higher pathway activity in a certain cancer type (or medium type or culture condition) was evaluated by a random permutation test. Cancer type labels (or media culture labels or culture condition labels) were randomly shuffled 5,000 times to simulate a null distribution of the pathway activity scores. We then statistically compared the shuffled pathway activity scores to the original, non-shuffled dataset and computed an empirical *P* value defined as the fraction of random pathway activity scores larger than *p_t,j_* (if *p_t,j_* > 1) or smaller than *p_t,j_* (if *p_t,j_* < 1) to determine if activity of a pathway is significantly higher or lower in a cancer type (or medium type or culture condition) than average. If a pathway activity was insignificant (i.e., *P* > 0.05), the activity score was assigned a value of 1. For the analysis of growth media and culture conditions, only those associated with at least 10 cell lines were used.

To determine the least and most variable metabolic pathways across cancer types, growth media, or culture conditions, we performed PCA on pathway activity scores. For each metabolic pathway, we calculated the sum of the absolute value of the loadings in the top 8 PCs (for cancer type analysis), the top 4 PCs (for media type analysis), or the top 2 PCs (for culture type analysis) as they accounted for at least 80% of the variance in data. To determine the sensitivity of analysis results to the variance threshold chosen (for capturing cancer type-associated heterogeneities), we repeated the analysis twice using 70% and 90% thresholds for data variance.

### Metabolomics data analysis and visualization across cancer cell lines

To systematically evaluate similarities and differences with respect to the abundance of metabolites across cell lines representing diverse cancer types, we used metabolomics data from the Cancer Cell Line Encyclopedia ^11^. The abundance of each of the 136 metabolites was z-scored across 875 cell lines representing 34 cancer types. UMAP clustering was performed in R using the z-scored data and the following parameters: nearest neighbor (n_neighbors) = 20, minimum distance (min_dist) = 0.3, and distance metric (metric) = Pearson. To assess variability across cancer types, we identified the median abundance for each metabolite for cancer types that had at least 5 cell lines profiled (29 cancer types). For each metabolite, we computed the interquartile range (IQR) across cancer type median values. We ordered metabolites based on their IQR values. We identified the top 50% variable metabolites using a 50-percentile threshold on IQR values. To assess the extent of cancer type-associated pathway variability based on metabolomics data, we used enrichment ratio derived from pathway enrichment analysis of the top 50% variable metabolites using the KEGG database in Metaboanalyst (https://www.metaboanalyst.ca/) ^30^.

### Analysis of cell line-to-cell line heterogeneity in metabolic pathways

To determine the extent of cell line-to-cell line heterogeneity in each cancer type, we used a three-step procedure adapted from Xiao *et al* ^29^. Frist, we performed PCA on z-scored metabolic gene expression data within each cancer type. We computed metabolic gene variability scores as the absolute sum of the PCA loading values across the top PCs accounting for at least 80% of the variance in data. To identify which metabolic pathways were most variable across cell lines within each cancer type, we applied pre-ranked gene set enrichment analysis (GSEA) to the ranked lists of metabolic gene variability scores. Scores were ranked in descending order and ran in pre-rank GSEA using the *fgseaSimple* function in the *fgsea* package in R (1.20.0). GSEA was performed against the same 85 KEGG metabolic pathways. A cancer type was determined to be significantly heterogeneous for a metabolic pathway if the GSEA generated FDR ≤ 0.05. If a cancer type was significantly heterogeneous for a specific metabolic pathway, we calculated metabolic state scores (e.g., Oxphos state scores) across the cell lines within the cancer type. Metabolic state scores were calculated as mean z-score values across the genes found within the specific metabolic pathway.

To test the generalizability of the findings regarding cell line-to-cell line heterogeneity within each cancer type, we repeated the analysis using metabolic gene expression data for an independent set of cell lines from the Sanger Cell Lines Project (v99) ^33^. Cancer types from the Sanger dataset were selected if 5 or more cell lines were RNA-sequenced, resulting in 891 cell lines spanning 33 cancer types. Z-scored expression of 1,497 metabolic genes was used as provided by the Sanger Institute. To assess the overall consistency of metabolic pathway activities (inferred based on transcriptomics data) between CCLE and Sanger datasets, we performed Pearson’s correlation analysis for each gene across the 658 cell lines shared between the two datasets. Genes were considered to show substantial differences between CCLE and Sanger datasets if their correlation coefficient (*r*) was <0.5. These genes were mapped back to the KEGG database. Metabolic pathways were considered to show substantial differences between CCLE and Sanger datasets based on the percentage of their constituent genes with *r* < 0.5. Quantitative analysis of cell line-to-cell line heterogeneity on Sanger cell lines (including PCA followed by GSEA) was performed using the same approach as described above.

### Analysis of cell-to-cell heterogeneity in Oxphos state across cell lines

To analyze cell-to-cell heterogeneity in Oxphos state across cell lines, single-cell RNA-sequencing data from Kinker *et al* ^31^ was used. Single-cell Oxphos state scores were computed as the average of counts per million (CPM) values for the expression of 101 Oxphos genes and reported for representative Oxphos^High^ and Oxphos^Low^ colorectal adenocarcinoma and pancreatic adenocarcinoma cell lines. An arbitrary CPM threshold of 200 was used to determine the percentage of Oxphos^High^ and Oxphos^Low^ single cells for each cell line.

### Feature selection via elastic net regularization to reveal pathway dependencies associated with Oxphos state

To systematically identify gene vulnerabilities associated with Oxphos state, we performed feature selection and multivariate modeling using DepMap gene dependency scores, representing the effect of genome-wide knockout effects on cell viability. A dependency score of 0 indicates no inhibitory effect (corresponding to a non-essential gene), and 1 indicates complete inhibitory effect (corresponding to an essential gene) ^34^. We first calculated Oxphos state scores across the cell lines, defined as the average of z-scores for the expression levels of 113 Oxphos genes. To amplify the impact of Oxphos^High^ and Oxphos^Low^ groups of cell lines in feature selection, we ordered cell lines based on their Oxphos state scores and included only the top and bottom 33 percentiles, resulting in 328 cell lines. As the first step of feature selection, we removed genes with dependency scores that varied minimally across Oxphos^High^ and Oxphos^Low^ cell lines. To this end, gene dependency score interquartile range (IQR) values were calculated across cell lines and genes with IQR < 0.09 were removed, narrowing down the list of genes from 17,202 to 3,624. We then used elastic net regularization, which integrates the penalty functions of least absolute shrinkage and selection operator (LASSO; L1 penalty) and ridge regression (L2 penalty), to further reduce the number of features and select a subset of the remaining genes based on their association with the Oxphos state score. Elastic net was run in R using the *cv.glmnet* function in the *glmnet* package (4.1-7). We first optimized the alpha (α), the parameter that controls the weight of L1 and L2 penalties, across the 296 cell lines in the training dataset. We held out a random set of 32 cell lines with variable Oxphos state scores as the test dataset for independent validation. 150 elastic net iterations (using 10-fold cross validation) were run with α ranging from 0.1 to 0.9 for a total of 1,350 models. We reported the smallest mean squared error (MSE) associated with each model based on the minimum lambda (*lambda.min* or λ_min_). Optimized elastic net α was determined based on the smallest reported median MSE. We selected genes that appeared in at least 50% of the 150 optimized elastic net iterations. The list of 200 selected genes with positive and negative mean coefficients were used in Enrichr (https://maayanlab.cloud/Enrichr/) ^66^ to determine pathways and cellular components associated with Oxphos^High^ and Oxphos^Low^ state dependencies, respectively.

To evaluate the overall generalizability of Oxphos vulnerabilities captured in DepMap data, we performed an independent but similar analysis (including feature selection followed by enrichment analysis) using CRISPR knockout scores provided in the Sanger dataset ^35^. A knockout score of 0 indicates no inhibitory effect and 1 indicates complete inhibitory effect. We first calculated Oxphos state scores across 225 cell lines found within the 15 Oxphos variable cancer types within the Sanger dataset. Oxphos state scores were defined as the average of z-scores for the expression levels of 100 Oxphos genes as reported in the Sanger dataset. To amplify the impact of Oxphos^High^ and Oxphos^Low^ groups of cell lines in feature selection, we ordered cell lines based on their Oxphos state scores and included only the top and bottom 33 percentiles, resulting in 148 cell lines. As the first step of feature selection, we removed genes in which their knockout scores were 0 or 1 across all 148 cell lines, narrowing down the list of gene knockouts from 17,995 to 6,721. Elastic net alpha (α) optimization and modeling and gene set enrichment were performed as described above. We optimized α across the 148 cell lines, α = 0.1. We identified 192 gene knockouts associated with Oxphos^High^ and Oxphos^Low^ cell lines.

### Partial least squares regression modeling to uncover most predictive gene vulnerabilities associated with Oxphos state

To quantify the relative importance of 200 gene dependencies (identified by elastic net regularization) in predicting the Oxphos state of each CCLE cell line, we built a partial least-square regression (PLSR) model ^36,37^ using the built-in MATLAB function *plsregress*. To evaluate the predictability of the linear relationship between the input (i.e., gene dependency values) and output variables (i.e., Oxphos state scores), we used leave-one-out cross-validation. The goodness of fit for each model was calculated using R^2^. Prediction accuracy was evaluated by Q^2^ and pairwise Pearson’s correlations between the measured and predicted Oxphos state scores. To independently validate the model, we also used the test set of 32 cell lines that were not included in either elastic net regularization or PLSR training steps. For the assessment of relative variable importance in the PLSR model, the information content of each variable was assessed by its variable importance in the projection (VIP) score ^70^. Based on |VIP| ≥ 1, we identified 64 genes whose dependency scores appeared to be the most significant predictors of the Oxphos state.

To further narrow down the list of genes (based on significance) for the next steps of analysis, we used a combination of correlation and permutation testing. Oxphos state scores for all 495 cell lines were correlated with their associated gene dependency scores for each of the 64 genes using Pearson’s correlation (R built-in function *cor.test*). Correlation *P* values were adjusted by calculating FDR values using R’s built-in function *p.adjust*. In the permutation test, we assessed the gene knockout effect in Oxphos^High^ versus Oxphos^Low^ cell lines (164 cell lines per group) by calculating the difference between the states’ median gene dependency scores. A positive difference indicates Oxphos^High^ cell lines are more sensitive to the gene knockout (i.e., Oxphos^High^ vulnerability) while a negative difference indicates Oxphos^Low^ cell lines are more sensitive to the gene knockout (i.e., Oxphos^Low^ vulnerability). Statistical significance of gene knockout effect was evaluated by a random permutation test. Oxphos states (i.e., Oxphos^High^, Oxphos^Low^) were randomly shuffled 8,000 times to simulate a null distribution of the differences between the states’ median gene dependency. We then statistically compared the shuffled differences to the original, non-shuffled dataset and computed an empirical *P* value to determine if gene knockout effect is a significant Oxphos^High^ or Oxphos^Low^ vulnerability. The overlap between correlation and permutation analyses found 53/64 gene knockouts to be significant (*P* ≤ 0.05 for permutation test and FDR ≤ 0.05 for Pearson correlation). Among these genes, we focused on those whose median dependencies between subgroups of Oxphos^High^ and Oxphos^Low^ cell lines were larger than 0.15. This led to a final list of 12 gene vulnerabilities strongly associated with the Oxphos^High^ state and 15 gene vulnerabilities strongly associated with the Oxphos^Low^ state.

### Analysis of statistical associations between Oxphos state, driver mutations and tissue context

To determine whether any of the Oxphos state-associated gene vulnerabilities were enriched more significantly in cell lines originating from specific tissue types or those carrying specific driver mutations, we compared median gene dependency between Oxphos^High^ and Oxphos^Low^ subgroups among cell lines grouped based on cancer type or occurrence of the top 25 commonly mutated driver genes ^39^. Among these genes, we focused on the top eleven, which carried mutations in at least 15 tested cell lines, including TP53, KRAS, PIK3CA, BRAF, APF, ARID1A, PTEN, NF1, FAT1, KMT2D, CREBBP. To assess the statistical significance of the observed differences in gene dependency, we performed permutation testing by randomly shuffling (8,000 times for each cancer type; 4,000 times for each mutation) the Oxphos state labels (High and Low) across cell lines participating in each comparison and computing permutation *P* values. We then used *P* ≤ 0.01 as the significance cutoff to identify mutations and tissue types associated with enhanced Oxphos state-specific gene dependencies relative to pan-cancer analysis.

### Analysis of Oxphos state correlation with cell line growth rate (doubling time)

To assess whether the Oxphos state of a cell line is associated with its growth rate (doubling time), we used a two-sample, two-sided *t* test using R’s built in *t.test* function to compare the doubling time of Oxphos^High^ and Oxphos^Low^ cell lines. Doubling time information was downloaded from Cellosaurus ^65^.

### Celligner analysis to integrate and align metabolic gene expression data across cell lines and patient-derived tumors

To assess the similarities and differences in metabolic gene expression data across CCLE cell lines and patient-derived tumors included in TCGA, TARGET, and Treehouse databases ^27,28^, we used aligned RNA-sequencing data from Celligner ^26^. The twenty common cancer types that represent both cell lines and tumors included glioma, osteosarcoma, acute lymphoblastic leukemia, embryonal rhabdomyosarcoma, alveolar rhabdomyosarcoma, acute myeloid leukemia, Ewing sarcoma, synovial sarcoma, ovarian serous cystadenocarcinoma, hepatocellular carcinoma, thyroid carcinoma, colon adenocarcinoma, non-small cell lung cancer, melanoma, renal cell carcinoma, prostate adenocarcinoma, breast, esophageal carcinoma, pancreas adenocarcinoma, and uveal melanoma. UMAP clustering in R was performed across 623 cell lines and 7,258 tumors based on the expression of 1,604 metabolic genes using the aligned RNA-sequencing data and the following parameters: nearest neighbor (n_neighbors) = 20, minimum distance (min_dist) = 0.5, and distance metric (metric) = Pearson. Percentage of cell lines of each cancer type grouped with their corresponding tumor type of origin was calculated using functions developed by Warren *et al* ^26^, including *calc_tumor_CL_cor*, *get_cell_line_tumor_class*, *cell_line_tumor_class*, and *cell_line_tumor_class_plot* in R (https://github.com/broadinstitute/Celligner_ms; retrieved 17 January 2024).

### Bioinformatics analysis on patient-derived glioma tumors

Clinical history, mutation profiles, and gene expression correlation were accessed using the cBioPortal for Cancer Genomics (http://cbioportal.org/) ^71^. Diffuse glioma patient samples from the GLASS Consortium were used for analysis ^47^. Diffuse glioma patients were classified based on their PTEN mutation status, resulting in 64 patients with PTEN mutations (PTEN^Mut^) and 158 patients with wildtype PTEN (PTEN^WT^). Of these patients, 57 PTEN^Mut^ and 150 PTEN^WT^ had survival data. Overall survival of PTEN^Mut^ versus PTEN^WT^ patients were compared using two-sided *P* value computed by log-rank (Mantel–Cox) test. To infer how PTEN expression might be associated with glioma tumor state, we performed transcriptome-wide co-expression analysis across 79 tumor samples for which RNA-sequencing data were available. We used Spearman’s correlation analysis to rank transcripts based on the association of their abundance with PTEN expression. *P* values were then corrected using Benjamini-Hochberg FDR correction. Genes that were negatively or positively correlated with PTEN expression and had FDR ≤ 0.05 were selected for gene set enrichment analysis using Enrichr.

### Single-cell RNA sequencing analysis of glioma tumors

To test the association of PTEN loss with mitochondrial electron transport chain (ETC) components at a single-cell level, we analyzed a published single-cell RNA-sequencing dataset of genetically profiled patient-derived glioblastoma tumors ^48^. Malignant cells were classified based on the PTEN mutation status, resulting in 10,268 PTEN-mutant cells found in 6 patients and 23,229 PTEN-wildtype cells found in 20 patients. For gene expression analysis, we used the log-normalized gene expression data, as reported by Ruiz-Moreno *et al* ^48^, for computing mean gene expression scores. Mean gene expression scores were calculated for every single cell using the follow gene sets: Respiratory Chain Complex I (GO:0045271), Respiratory Chain Complex II (GO:0045273), Respiratory Chain Complex III (GO:0005750), Respiratory Chain Complex IV (GO:0045277), and ATP synthase (GO:0045259). Significance of gene set expression between PTEN-mutant and PTEN-wildtype cells was determined using a one-sided, permutation test (5,000 permutations per test), hypothesizing that PTEN-mutant cells are higher in gene expression compared to PTEN-wildtype. The permutation test MATLAB function was accessed via MathWorks File Exchange (https://github.com/lrkrol/permutationTest; retrieved 13 September 2023).

### Drug sensitivity analysis using the Cancer Therapeutics Response Portal (CTRP)

To test the impact of damaging mutations in PTEN on Oxphos state-specific dependencies, we used the Cancer Therapeutics Response Portal (CTRP) data to analyze the sensitivity of 799 genetically characterized cancer cell lines to 545 small-molecule probes and drugs ^40^. We used transcriptomics data to define groups of Oxphos^Low^ and Oxphos^High^ cell lines based on whether their Oxphos state scores were ranked within the top or bottom 33 percentiles. We then compared the median sensitivity of Oxphos^Low^ and Oxphos^High^ cell lines to each tested small molecule based on the area of the dose-response curve (AUC) measurements. Within the Oxphos^High^ group, we also compared the median sensitivity of PTEN-wildtype (PTEN^WT^) and PTEN-mutated (PTEN^Mut^) subgroups. Significance of comparison was determined using one-sided Wilcoxon rank sum test by MATLAB function *ranksum*.

### Hierarchical clustering

Unsupervised hierarchical clustering of pathway activity scores and *P* values from pre-rank GSEA were carried out in R using the *pheatmap* package (1.0.12). Clustering was performed using the *pheatmap* function using default settings with Euclidean distance metric.

### Overexpression of PTEN in intrinsically PTEN^Null^ U251MG glioblastoma cells

To generate stable PTEN-overexpressing (PTEN^OE^) cells from the intrinsically PTEN^Null^ U251MG glioblastoma cell line, cells were seeded in antibiotic-free growth media in 6-well polystyrene tissue culture treated plates at a density of 100,000 cells/well. Two days after seeding, cells were infected with Precision LentiORF PTEN or Precision LentiORF positive control (RFP control) viral particles (purchased from Horizon Discovery) at a multiplicity of infection (MOI) of 0.3 in antibiotic and serum-free growth media. Each well received 1 mL of the appropriate viral particles in antibiotic and serum-free growth media and 10 μg/mL polybrene. The plate was centrifuged at 2,500 rpm for 30 min at 37°C. Five hours post-transduction, 3 mL of antibiotic-free growth media was added to each well. At 48 hours post-transduction, media was replaced with growth media supplemented with selection antibiotic, Blasticidin S Hydrochloride, at a concentration of 5 μg/mL to select positive clones. This concentration was chosen via dose-response analysis of Blasticidin S Hydrochloride in U251MG cells, demonstrating its effectiveness in killing non-transduced cells within 10 days.

### Drug sensitivity analysis via dose-response measurements of normalized growth rate

Growth rate inhibition assays with ETC inhibitors were performed in 96-well clear-bottom, black-walled plates excluding the outer wells which were filled with media. Viable cells were counted using a TC20 automated cell counter or a hemocytometer. For primary mouse glioma cells, the 96-well plates were coated with Poly-D-lysine and cells were seeded at a density of 10,000 per well in 200 μL of the above-mentioned media. For experiments with the U251MG cell line, cells transduced with lentiviral PTEN ORF (PTEN^OE^) and RFP control (Control) were seeded in 200 μL of full growth media at a density of 3,000 and 1,500 cells per well, respectively. The small-molecule inhibitors IACS-010759, oligomycin A, antimycin A, and Gboxin were dissolved in DMSO at a stock concentration of 10 mmol/L, and metformin was dissolved in cell-culture grade sterile water at 100 mmol/L.

Cells were treated the day after seeding with compounds at reported concentrations or with vehicle control (DMSO or water). DMSO and DMSO-based compounds were dispensed using a D300e Digital Dispenser for a final DMSO concentration of less than 0.1%. Water and water-based compounds were added manually for a final water concentration of less than 10%. At designated timepoints after drug treatment, including *t* = 0 h (i.e., no drug treatment), 96 h and 144 h (or 168 h), Hoechst 33342 in PBS was added to each well to a concentration of 1 μg/mL and plates were returned to the incubator. After an hour of incubation in Hoechst, cells were fixed with 4% paraformaldehyde (PFA) for 30 minutes at room temperature. Plates were then washed 4 times with PBS and imaged on the Operetta CLS High-Content Imaging System. For experiments with mouse primary glioma cells, a 20x objective was used and 25 sites were imaged per well. For experiments with U251MG cells, a 10x objective was used and a total of nine sites were imaged per well. Background subtraction was performed with Fiji (2.9.0) software. Image segmentation was performed with CellProfiler (4.2.5). Data were analyzed using MATLAB 2023a.

Drug sensitivity across cell lines was compared based on their normalized growth rate measured in the presence of each compound (at different doses) relative to vehicle control as previously described ^72^. To this end, net growth rate under each treatment condition was first calculated according to the following equation:

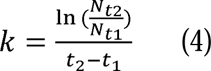

where *N_t1_* and *N_t2_* represent the number of cells measured at timepoints *t* = *t_1_* and *t* = *t_2_*, respectively, and *k* describes the net growth rate of cells during the period between *t_1_* and *t_2_*. The net growth rates were then averaged across multiple, consecutive time intervals (e.g., 0-96 hours and 96-144 hours) to determine the mean net growth rate for each treatment condition. To compare drug sensitivity among different cell lines, we normalized the mean net growth rate measured for drug-treated cells (*k_drug_*) to that measured for vehicle (DMSO)-treated cells (*k_DMSO_*) to calculate the normalized growth rate:

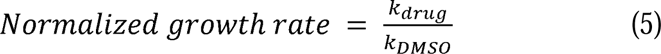

### Western blots

Cultured cells were transferred to ice, washed with ice-cold PBS, and lysed with ice-cold RIPA buffer consisting of 25 mmol/L Tris-HCl (pH 7.4), 150 mmol/L NaCl, 1% TritonX-100, 0.5% sodium deoxycholate, and 0.1% SDS supplemented with protease cocktail including 5 μg/mL leupeptin, 2 μg/mL aprotinin, 2 μg/mL pepstatin A and 0.2 mM AEBSF for mouse cell experiments or 150 mM NaCl, 1.0% IGEPAL® CA-630, 0.5% sodium deoxycholate, 0.1% SDS, and 50 mM Tris (pH 8.0) supplemented with 1 mmol/L Na_3_VO_4_, 15 mmol/L NaF, cOmplete ULTRA ethylene diamine tetraacetic acid (EDTA)–free protease inhibitors and PhosSTOP for human cell line experiments. Protein samples of cleared lysates were measured using BCA assay. Lysates were adjusted to equal protein concentrations for each sample within a blot (36-42 μg), 4x NuPage LDS sample buffer supplemented with 1 mL of 1 M DTT was added, and samples were heated for 10 minutes at 70°C. Samples were then loaded on NuPAGE 10% Bis-Tris gels. Western blots were performed using iBlot 2 Gel Transfer Stacks PVDF system. Membranes were blocked with Intercept (TBS) blocking buffer at room temperature for 1 hour and then incubated with primary antibodies (1:1,000 dilution for PTEN; 1:800 for β-tubulin) in Intercept blocking buffer overnight at 4°C. Secondary antibodies (anti-rabbit IRDye 800CW and anti-mouse IRDye 680RD) were used at 1:15,000 dilution in Intercept blocking buffer. Membranes were scanned on an Odyssey CLx scanner with 700 and 800 nm channels set to automatic intensity at 169 μm resolution. Blot images were processed on Odyssey application software (3.0.21).

### Quantification and statistical analysis

All boxplots highlight the median, lower, and upper quartiles. All violin plots highlight the median values. Whiskers in boxplots indicate 1.5 times interquartile ranges. Sample size (i.e., number of cell lines, cells, or replicates) are indicated in the figure legend. The significance of pairwise correlations were evaluated based on *P* values associated with the corresponding two-sided Pearson’s correlation analysis or Spearman’s correlation analysis. Statistical significance of metabolic pathway activities across cancer types, growth media, and culture type were determined based on permutation testing. The significance of knockout effects in Oxphos^High^ and Oxphos^Low^ cell lines across different mutations and tissue types was determined based on permutation testing. Statistical significance of mitochondrial gene expression in PTEN^Mut^ and PTEN^WT^ glioma single cells was determined based on permutation testing. Drug sensitivity analysis of data from the Cancer Therapeutics Response Portal (CTRP) for PTEN^Mut^ and PTEN^WT^ cell lines was performed using one-sided Wilcoxon rank sum test. Two-sided, two-sample *t* test was used to analyze the difference in doubling time between Oxphos^High^ and Oxphos^Low^ cell lines. Statistical significance of the effect of ETC inhibitors on PTEN^Null^ vs. PTEN^WT^ or PTEN^OE^ vs. Control cell lines was determined by two-way ANOVA.

## Acknowledgments

We thank D. Kashatus, B. Paudel, Papin laboratory, members of the Fallahi-Sichani laboratory and the Systems Analysis of Stress-adapted Cancer Organelles (SASCO) Center at UVA for helpful discussion and suggestions. This work was supported by NIH grants R35-GM133404, R01-CA249229, U54-CA274499, R21-NS125479, T32-GM145443, University of Michigan Rackham Merit Fellowship (to C.A.) and UVA Engineering Distinguished Fellowship (to A.D.K.).

## Author Contributions

C.A. and M.F.-S. conceived and designed the study. C.A. performed the computational modeling work. A.D.K. and Y.J. performed the experiments. A.D.K. analyzed the experimental data. C.A., A.D.K., Y.J., H.Z., and M.F.-S wrote and edited the manuscript. M.F.-S. supervised the work.

## State of Competing Interests

The authors declare that they have no competing interests.

## References

1. Finley, L.W.S. (2023). What is cancer metabolism? Cell 186, 1670–1688. 10.1016/j.cell.2023.01.038.

2. Stine, Z.E., Schug, Z.T., Salvino, J.M., and Dang, C.V. (2022). Targeting cancer metabolism in the era of precision oncology. Nat Rev Drug Discov 21, 141–162. 10.1038/s41573-021-00339-6.

3. Bi, J., Wu, S., Zhang, W., and Mischel, P.S. (2018). Targeting cancer’s metabolic co-dependencies: A landscape shaped by genotype and tissue context. Biochim Biophys Acta Rev Cancer 1870, 76–87. 10.1016/j.bbcan.2018.05.002.

4. Fendt, S.-M., Frezza, C., and Erez, A. (2020). Targeting Metabolic Plasticity and Flexibility Dynamics for Cancer Therapy. Cancer Discovery 10, 1797–1807. 10.1158/2159-8290.CD-20-0844.

5. Kondo, H., Ratcliffe, C.D.H., Hooper, S., Ellis, J., MacRae, J.I., Hennequart, M., Dunsby, C.W., Anderson, K.I., and Sahai, E. (2021). Single-cell resolved imaging reveals intra-tumor heterogeneity in glycolysis, transitions between metabolic states, and their regulatory mechanisms. Cell Reports 34, 108750. 10.1016/j.celrep.2021.108750.

6. Rohle, D., Popovici-Muller, J., Palaskas, N., Turcan, S., Grommes, C., Campos, C., Tsoi, J., Clark, O., Oldrini, B., Komisopoulou, E., et al. (2013). An inhibitor of mutant IDH1 delays growth and promotes differentiation of glioma cells. Science 340, 626–630. 10.1126/science.1236062.

7. Yen, K., Travins, J., Wang, F., David, M.D., Artin, E., Straley, K., Padyana, A., Gross, S., DeLaBarre, B., Tobin, E., et al. (2017). AG-221, a First-in-Class Therapy Targeting Acute Myeloid Leukemia Harboring Oncogenic IDH2 Mutations. Cancer Discov 7, 478–493. 10.1158/2159-8290.CD-16-1034.

8. Kim, J., Hu, Z., Cai, L., Li, K., Choi, E., Faubert, B., Bezwada, D., Rodriguez-Canales, J., Villalobos, P., Lin, Y.-F., et al. (2017). CPS1 maintains pyrimidine pools and DNA synthesis in KRAS/LKB1-mutant lung cancer cells. Nature 546, 168–172. 10.1038/nature22359.

9. Nwosu, Z.C., Ward, M.H., Sajjakulnukit, P., Poudel, P., Ragulan, C., Kasperek, S., Radyk, M., Sutton, D., Menjivar, R.E., Andren, A., et al. (2023). Uridine-derived ribose fuels glucose-restricted pancreatic cancer. Nature 618, 151–158. 10.1038/s41586-023-06073-w.

10. Gwynne, W.D., Suk, Y., Custers, S., Mikolajewicz, N., Chan, J.K., Zador, Z., Chafe, S.C., Zhai, K., Escudero, L., Zhang, C., et al. (2022). Cancer-selective metabolic vulnerabilities in MYC-amplified medulloblastoma. Cancer Cell 40, 1488–1502.e7. 10.1016/j.ccell.2022.10.009.

11. Li, H., Ning, S., Ghandi, M., Kryukov, G.V., Gopal, S., Deik, A., Souza, A., Pierce, K., Keskula, P., Hernandez, D., et al. (2019). The landscape of cancer cell line metabolism. Nat Med 25, 850–860. 10.1038/s41591-019-0404-8.

12. Ghandi, M., Huang, F.W., Jané-Valbuena, J., Kryukov, G.V., Lo, C.C., McDonald, E.R., Barretina, J., Gelfand, E.T., Bielski, C.M., Li, H., et al. (2019). Next-generation characterization of the Cancer Cell Line Encyclopedia. Nature 569, 503–508. 10.1038/s41586-019-1186-3.

13. Joly, J.H., Chew, B.T.L., and Graham, N.A. (2021). The landscape of metabolic pathway dependencies in cancer cell lines. PLoS Comput Biol 17, e1008942. 10.1371/journal.pcbi.1008942.

14. Lagziel, S., Lee, W.D., and Shlomi, T. (2019). Inferring cancer dependencies on metabolic genes from large-scale genetic screens. BMC Biol 17, 37. 10.1186/s12915-019-0654-4.

15. Cherkaoui, S., Durot, S., Bradley, J., Critchlow, S., Dubuis, S., Masiero, M.M., Wegmann, R., Snijder, B., Othman, A., Bendtsen, C., et al. (2022). A functional analysis of 180 cancer cell lines reveals conserved intrinsic metabolic programs. Molecular Systems Biology 18, e11033. 10.15252/msb.202211033.

16. Shorthouse, D., Bradley, J., Critchlow, S.E., Bendtsen, C., and Hall, B.A. (2022). Heterogeneity of the cancer cell line metabolic landscape. Molecular Systems Biology 18, e11006. 10.15252/msb.202211006.

17. Pemovska, T., Bigenzahn, J.W., Srndic, I., Lercher, A., Bergthaler, A., César-Razquin, A., Kartnig, F., Kornauth, C., Valent, P., Staber, P.B., et al. (2021). Metabolic drug survey highlights cancer cell dependencies and vulnerabilities. Nat Commun 12, 7190. 10.1038/s41467-021-27329-x.

18. Leeuwenburgh, V.C., Urzúa-Traslaviña, C.G., Bhattacharya, A., Walvoort, M.T.C., Jalving, M., de Jong, S., and Fehrmann, R.S.N. (2021). Robust metabolic transcriptional components in 34,494 patient-derived cancer-related samples and cell lines. Cancer Metab 9, 35. 10.1186/s40170-021-00272-7.

19. Yang, J., Griffin, A., Qiang, Z., and Ren, J. (2022). Organelle-targeted therapies: a comprehensive review on system design for enabling precision oncology. Signal Transduct Target Ther 7, 379. 10.1038/s41392-022-01243-0.

20. Benedetti, E., Liu, E.M., Tang, C., Kuo, F., Buyukozkan, M., Park, T., Park, J., Correa, F., Hakimi, A.A., Intlekofer, A.M., et al. (2023). A multimodal atlas of tumour metabolism reveals the architecture of gene-metabolite covariation. Nat Metab 5, 1029–1044. 10.1038/s42255-023-00817-8.

21. Campit, S.E., Bhowmick, R., Lu, T., Saoji, A.V., Jin, R., Robida, A.M., and Chandrasekaran, S. (2023). Data-Driven Screening to Infer Metabolic Modulators of the Cancer Epigenome (Systems Biology) 10.1101/2023.02.27.530260.

22. Sessions, D.T., Kim, K.-B., Kashatus, J.A., Churchill, N., Park, K.-S., Mayo, M.W., Sesaki, H., and Kashatus, D.F. (2022). Opa1 and Drp1 reciprocally regulate cristae morphology, ETC function, and NAD+ regeneration in KRas-mutant lung adenocarcinoma. Cell Rep 41, 111818. 10.1016/j.celrep.2022.111818.

23. Barretina, J., Caponigro, G., Stransky, N., Venkatesan, K., Margolin, A.A., Kim, S., Wilson, C.J., Lehár, J., Kryukov, G.V., Sonkin, D., et al. (2012). The Cancer Cell Line Encyclopedia enables predictive modelling of anticancer drug sensitivity. Nature 483, 603–607. 10.1038/nature11003.

24. Becht, E., McInnes, L., Healy, J., Dutertre, C.-A., Kwok, I.W.H., Ng, L.G., Ginhoux, F., and Newell, E.W. (2019). Dimensionality reduction for visualizing single-cell data using UMAP. Nat Biotechnol 37, 38–44. 10.1038/nbt.4314.

25. Kanehisa, M., Sato, Y., Kawashima, M., Furumichi, M., and Tanabe, M. (2016). KEGG as a reference resource for gene and protein annotation. Nucleic Acids Research 44, D457–D462. 10.1093/nar/gkv1070.

26. Warren, A., Chen, Y., Jones, A., Shibue, T., Hahn, W.C., Boehm, J.S., Vazquez, F., Tsherniak, A., and McFarland, J.M. (2021). Global computational alignment of tumor and cell line transcriptional profiles. Nat Commun 12, 22. 10.1038/s41467-020-20294-x.

27. Weinstein, J.N., Collisson, E.A., Mills, G.B., Shaw, K.R.M., Ozenberger, B.A., Ellrott, K., Shmulevich, I., Sander, C., and Stuart, J.M. (2013). The Cancer Genome Atlas Pan-Cancer analysis project. Nat Genet 45, 1113–1120. 10.1038/ng.2764.

28. Goldman, M.J., Craft, B., Hastie, M., Repečka, K., McDade, F., Kamath, A., Banerjee, A., Luo, Y., Rogers, D., Brooks, A.N., et al. (2020). Visualizing and interpreting cancer genomics data via the Xena platform. Nat Biotechnol 38, 675–678. 10.1038/s41587-020-0546-8.

29. Xiao, Z., Dai, Z., and Locasale, J.W. (2019). Metabolic landscape of the tumor microenvironment at single cell resolution. Nat Commun 10, 3763. 10.1038/s41467-019-11738-0.

30. Xia, J., and Wishart, D.S. (2011). Web-based inference of biological patterns, functions and pathways from metabolomic data using MetaboAnalyst. Nat Protoc 6, 743–760. 10.1038/nprot.2011.319.

31. Kinker, G.S., Greenwald, A.C., Tal, R., Orlova, Z., Cuoco, M.S., McFarland, J.M., Warren, A., Rodman, C., Roth, J.A., Bender, S.A., et al. (2020). Pan-cancer single-cell RNA-seq identifies recurring programs of cellular heterogeneity. Nat Genet 52, 1208–1218. 10.1038/s41588-020-00726-6.

32. Ackermann, T., and Tardito, S. (2019). Cell Culture Medium Formulation and Its Implications in Cancer Metabolism. Trends Cancer 5, 329–332. 10.1016/j.trecan.2019.05.004.

33. Sondka, Z., Dhir, N.B., Carvalho-Silva, D., Jupe, S., Madhumita, McLaren, K., Starkey, M., Ward, S., Wilding, J., Ahmed M., et al. (2024). COSMIC: a curated database of somatic variants and clinical data for cancer. Nucleic Acids Research 52, D1210–D1217. 10.1093/nar/gkad986.

34. Dempster, J.M., Boyle, I., Vazquez, F., Root, D.E., Boehm, J.S., Hahn, W.C., Tsherniak, A., and McFarland, J.M. (2021). Chronos: a cell population dynamics model of CRISPR experiments that improves inference of gene fitness effects. Genome Biology 22, 343. 10.1186/s13059-021-02540-7.

35. Behan, F.M., Iorio, F., Picco, G., Gonçalves, E., Beaver, C.M., Migliardi, G., Santos, R., Rao, Y., Sassi, F., Pinnelli, M., et al. (2019). Prioritization of cancer therapeutic targets using CRISPR-Cas9 screens. Nature 568, 511–516. 10.1038/s41586-019-1103-9.

36. Gunn, B.M., Yu, W.-H., Karim, M.M., Brannan, J.M., Herbert, A.S., Wec, A.Z., Halfmann, P.J., Fusco, M.L., Schendel, S.L., Gangavarapu, K., et al. (2018). A Role for Fc Function in Therapeutic Monoclonal Antibody-Mediated Protection against Ebola Virus. Cell Host & Microbe 24, 221–233.e5. 10.1016/j.chom.2018.07.009.

37. Selva, K.J., van de Sandt, C.E., Lemke, M.M., Lee, C.Y., Shoffner, S.K., Chua, B.Y., Davis, S.K., Nguyen, T.H.O., Rowntree, L.C., Hensen, L., et al. (2021). Systems serology detects functionally distinct coronavirus antibody features in children and elderly. Nat Commun 12, 2037. 10.1038/s41467-021-22236-7.

38. Zhang, L., Cao, J., Dong, L., and Lin, H. (2020). TiPARP forms nuclear condensates to degrade HIF-1α and suppress tumorigenesis. Proc Natl Acad Sci U S A 117, 13447–13456. 10.1073/pnas.1921815117.

39. Mendiratta, G., Ke, E., Aziz, M., Liarakos, D., Tong, M., and Stites, E.C. (2021). Cancer gene mutation frequencies for the U.S. population. Nat Commun 12, 5961. 10.1038/s41467-021-26213-y.

40. Rees, M.G., Seashore-Ludlow, B., Cheah, J.H., Adams, D.J., Price, E.V., Gill, S., Javaid, S., Coletti, M.E., Jones, V.L., Bodycombe, N.E., et al. (2016). Correlating chemical sensitivity and basal gene expression reveals mechanism of action. Nat Chem Biol 12, 109–116. 10.1038/nchembio.1986.

41. Linnett, P.E., and Beechey, R.B. (1979). Inhibitors of the ATP synthetase systems. In Methods in Enzymology (Elsevier), pp. 472–518. 10.1016/0076-6879(79)55061-7.

42. Ulanovskaya, O.A., Janjic, J., Suzuki, M., Sabharwal, S.S., Schumacker, P.T., Kron, S.J., and Kozmin, S.A. (2008). Synthesis enables identification of the cellular target of leucascandrolide A and neopeltolide. Nat Chem Biol 4, 418–424. 10.1038/nchembio.94.

43. Jonsson, P., Lin, A.L., Young, R.J., DiStefano, N.M., Hyman, D.M., Li, B.T., Berger, M.F., Zehir, A., Ladanyi, M., Solit, D.B., et al. (2019). Genomic Correlates of Disease Progression and Treatment Response in Prospectively Characterized Gliomas. Clin Cancer Res 25, 5537–5547. 10.1158/1078-0432.CCR-19-0032.

44. Shi, Y., Lim, S.K., Liang, Q., Iyer, S.V., Wang, H.-Y., Wang, Z., Xie, X., Sun, D., Chen, Y.-J., Tabar, V., et al. (2019). Gboxin is an oxidative phosphorylation inhibitor that targets glioblastoma. Nature 567, 341–346. 10.1038/s41586-019-0993-x.

45. Molina, J.R., Sun, Y., Protopopova, M., Gera, S., Bandi, M., Bristow, C., McAfoos, T., Morlacchi, P., Ackroyd, J., Agip, A.-N.A., et al. (2018). An inhibitor of oxidative phosphorylation exploits cancer vulnerability. Nat Med 24, 1036–1046. 10.1038/s41591-018-0052-4.

46. Sesen, J., Dahan, P., Scotland, S.J., Saland, E., Dang, V.-T., Lemarié, A., Tyler, B.M., Brem, H., Toulas, C., Cohen-Jonathan Moyal, E., et al. (2015). Metformin inhibits growth of human glioblastoma cells and enhances therapeutic response. PLoS One 10, e0123721. 10.1371/journal.pone.0123721.

47. Barthel, F.P., Johnson, K.C., Varn, F.S., Moskalik, A.D., Tanner, G., Kocakavuk, E., Anderson, K.J., Abiola, O., Aldape, K., Alfaro, K.D., et al. (2019). Longitudinal molecular trajectories of diffuse glioma in adults. Nature 576, 112–120. 10.1038/s41586-019-1775-1.

48. Ruiz-Moreno, C., Salas, S.M., Samuelsson, E., Brandner, S., Kranendonk, M.E.G., Nilsson, M., and Stunnenberg, H.G. (2022). Harmonized single-cell landscape, intercellular crosstalk and tumor architecture of glioblastoma (Cancer Biology) 10.1101/2022.08.27.505439.

49. Ledur, P.F., Liu, C., He, H., Harris, A.R., Minussi, D.C., Zhou, H.-Y., Shaffrey, M.E., Asthagiri, A., Lopes, M.B.S., Schiff, D., et al. (2016). Culture conditions tailored to the cell of origin are critical for maintaining native properties and tumorigenicity of glioma cells. Neuro Oncol 18, 1413–1424. 10.1093/neuonc/now062.

50. Mahendralingam, M.J., Kim, H., McCloskey, C.W., Aliar, K., Casey, A.E., Tharmapalan, P., Pellacani, D., Ignatchenko, V., Garcia-Valero, M., Palomero, L., et al. (2021). Mammary epithelial cells have lineage-rooted metabolic identities. Nat Metab 3, 665–681. 10.1038/s42255-021-00388-6.

51. Du, J., Su, Y., Qian, C., Yuan, D., Miao, K., Lee, D., Ng, A.H.C., Wijker, R.S., Ribas, A., Levine, R.D., et al. (2020). Raman-guided subcellular pharmaco-metabolomics for metastatic melanoma cells. Nat Commun 11, 4830. 10.1038/s41467-020-18376-x.

52. Han, M., Bushong, E.A., Segawa, M., Tiard, A., Wong, A., Brady, M.R., Momcilovic, M., Wolf, D.M., Zhang, R., Petcherski, A., et al. (2023). Spatial mapping of mitochondrial networks and bioenergetics in lung cancer. Nature 615, 712–719. 10.1038/s41586-023-05793-3.

53. Cogliati, S., Frezza, C., Soriano, M.E., Varanita, T., Quintana-Cabrera, R., Corrado, M., Cipolat, S., Costa, V., Casarin, A., Gomes, L.C., et al. (2013). Mitochondrial cristae shape determines respiratory chain supercomplexes assembly and respiratory efficiency. Cell 155, 160–171. 10.1016/j.cell.2013.08.032.

54. Anderson, G.R., Wardell, S.E., Cakir, M., Yip, C., Ahn, Y.-R., Ali, M., Yllanes, A.P., Chao, C.A., McDonnell, D.P., and Wood, K.C. (2018). Dysregulation of mitochondrial dynamics proteins are a targetable feature of human tumors. Nat Commun 9, 1677. 10.1038/s41467-018-04033-x.

55. Naguib, A., Mathew, G., Reczek, C.R., Watrud, K., Ambrico, A., Herzka, T., Salas, I.C., Lee, M.F., El-Amine, N., Zheng, W., et al. (2018). Mitochondrial Complex I Inhibitors Expose a Vulnerability for Selective Killing of Pten-Null Cells. Cell Rep 23, 58–67. 10.1016/j.celrep.2018.03.032.

56. Yap, T.A., Daver, N., Mahendra, M., Zhang, J., Kamiya-Matsuoka, C., Meric-Bernstam, F., Kantarjian, H.M., Ravandi, F., Collins, M.E., Francesco, M.E.D., et al. (2023). Complex I inhibitor of oxidative phosphorylation in advanced solid tumors and acute myeloid leukemia: phase I trials. Nat Med 29, 115–126. 10.1038/s41591-022-02103-8.

57. Machado, N.D., Heather, L.C., Harris, A.L., and Higgins, G.S. (2023). Targeting mitochondrial oxidative phosphorylation: lessons, advantages, and opportunities. Br J Cancer 129, 897–899. 10.1038/s41416-023-02394-9.

58. Garofano, L., Migliozzi, S., Oh, Y.T., D’Angelo, F., Najac, R.D., Ko, A., Frangaj, B., Caruso, F.P., Yu, K., Yuan, J., et al. (2021). Pathway-based classification of glioblastoma uncovers a mitochondrial subtype with therapeutic vulnerabilities. Nat Cancer 2, 141–156. 10.1038/s43018-020-00159-4.

59. Sighel, D., Notarangelo, M., Aibara, S., Re, A., Ricci, G., Guida, M., Soldano, A., Adami, V., Ambrosini, C., Broso, F., et al. (2021). Inhibition of mitochondrial translation suppresses glioblastoma stem cell growth. Cell Rep 35, 109024. 10.1016/j.celrep.2021.109024.

60. Bi, J., Chowdhry, S., Wu, S., Zhang, W., Masui, K., and Mischel, P.S. (2020). Altered cellular metabolism in gliomas-an emerging landscape of actionable co-dependency targets. Nat Rev Cancer 20, 57–70. 10.1038/s41568-019-0226-5.

61. Hu, T., Allam, M., Cai, S., Henderson, W., Yueh, B., Garipcan, A., Ievlev, A.V., Afkarian, M., Beyaz, S., and Coskun, A.F. (2023). Single-cell spatial metabolomics with cell-type specific protein profiling for tissue systems biology. Nat Commun 14, 8260. 10.1038/s41467-023-43917-5.

62. Maan, K., Baghel, R., Dhariwal, S., Sharma, A., Bakhshi, R., and Rana, P. (2023). Metabolomics and transcriptomics based multi-omics integration reveals radiation-induced altered pathway networking and underlying mechanism. NPJ Syst Biol Appl 9, 42. 10.1038/s41540-023-00305-5.

63. Sun, C., Wang, A., Zhou, Y., Chen, P., Wang, X., Huang, J., Gao, J., Wang, X., Shu, L., Lu, J., et al. (2023). Spatially resolved multi-omics highlights cell-specific metabolic remodeling and interactions in gastric cancer. Nat Commun 14, 2692. 10.1038/s41467-023-38360-5.

64. Rees, M.G., Seashore-Ludlow, B., Cheah, J.H., Adams, D.J., Price, E.V., Gill, S., Javaid, S., Coletti, M.E., Jones, V.L., Bodycombe, N.E., et al. (2016). Correlating chemical sensitivity and basal gene expression reveals mechanism of action. Nat Chem Biol 12, 109–116. 10.1038/nchembio.1986.

65. Bairoch, A. (2018). The Cellosaurus, a Cell-Line Knowledge Resource. Journal of Biomolecular Techniques: JBT 29, 25. 10.7171/jbt.18-2902-002.

66. Kuleshov, M.V., Jones, M.R., Rouillard, A.D., Fernandez, N.F., Duan, Q., Wang, Z., Koplev, S., Jenkins, S.L., Jagodnik, K.M., Lachmann, A., et al. (2016). Enrichr: a comprehensive gene set enrichment analysis web server 2016 update. Nucleic Acids Res 44, W90–W97. 10.1093/nar/gkw377.

67. Stirling, D.R., Swain-Bowden, M.J., Lucas, A.M., Carpenter, A.E., Cimini, B.A., and Goodman, A. (2021). CellProfiler 4: improvements in speed, utility and usability. BMC Bioinformatics 22, 433. 10.1186/s12859-021-04344-9.

68. Liu, C., Sage, J.C., Miller, M.R., Verhaak, R.G.W., Hippenmeyer, S., Vogel, H., Foreman, O., Bronson, R.T., Nishiyama, A., Luo, L., et al. (2011). Mosaic analysis with double markers reveals tumor cell of origin in glioma. Cell 146, 209–221. 10.1016/j.cell.2011.06.014.

69. Lesche, R., Groszer, M., Gao, J., Wang, Y., Messing, A., Sun, H., Liu, X., and Wu, H. (2002). Cre/loxP-mediated inactivation of the murine Pten tumor suppressor gene. Genesis 32, 148–149. 10.1002/gene.10036.

70. Wold, S. (1994). Exponentially weighted moving principal components analysis and projections to latent structures. Chemometrics and Intelligent Laboratory Systems 23, 149–161. 10.1016/0169-7439(93)E0075-F.

71. Cerami, E., Gao, J., Dogrusoz, U., Gross, B.E., Sumer, S.O., Aksoy, B.A., Jacobsen, A., Byrne, C.J., Heuer, M.L., Larsson, E., et al. (2012). The cBio Cancer Genomics Portal: An Open Platform for Exploring Multidimensional Cancer Genomics Data. Cancer Discovery 2, 401–404. 10.1158/2159-8290.CD-12-0095.

72. Abecunas, C., Whitehead, C.E., Ziemke, E.K., Baumann, D.G., Frankowski-McGregor, C.L., Sebolt-Leopold, J.S., and Fallahi-Sichani, M. (2023). Loss of NF1 in Melanoma Confers Sensitivity to SYK Kinase Inhibition. Cancer Research 83, 316–331. 10.1158/0008-5472.CAN-22-0883.

