## Supplementary Figures for "Multivariate analysis of metabolic state vulnerabilities across diverse cancer contexts reveals synthetically lethal associations"

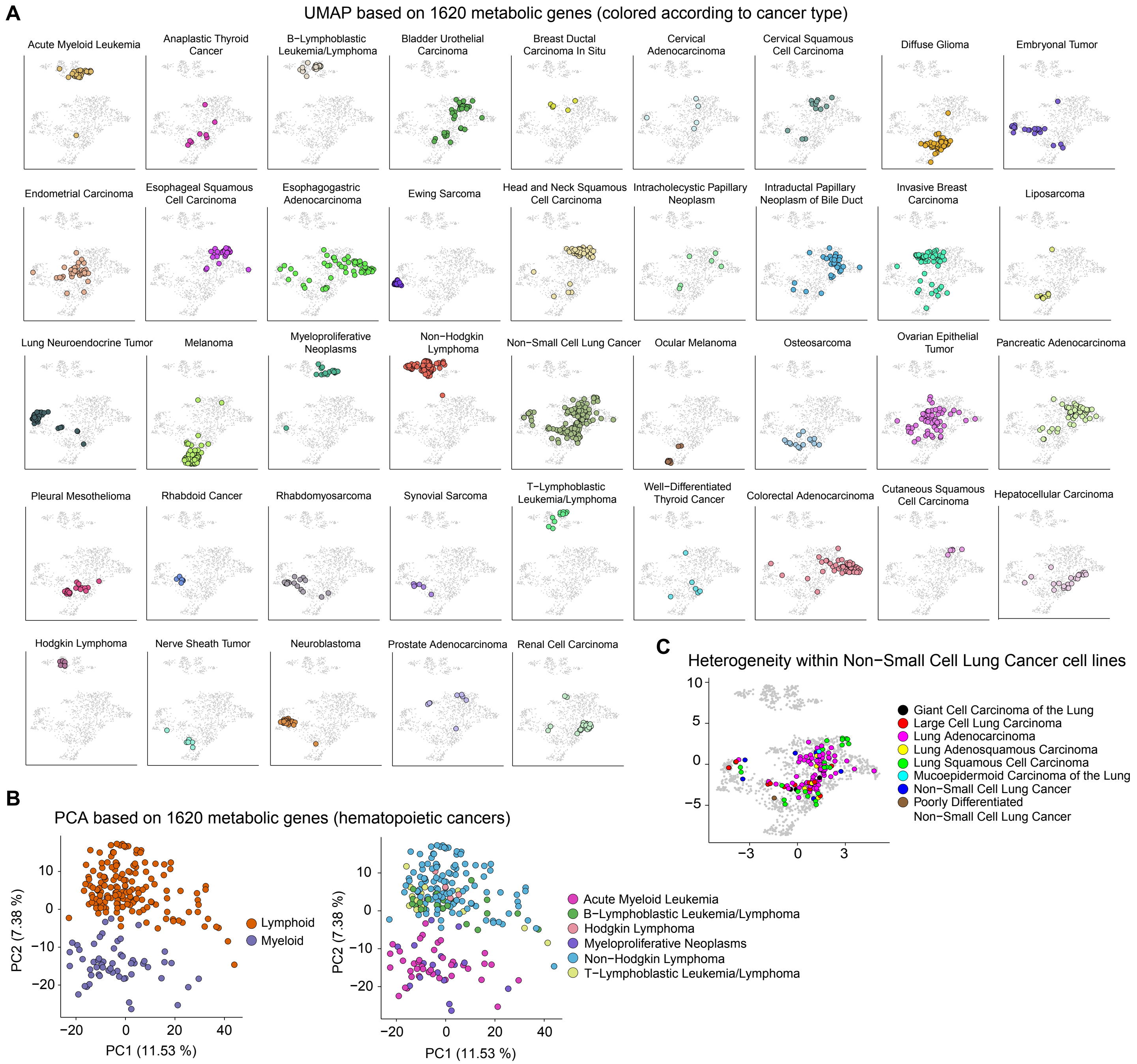


**Figure S1. UMAP analysis of metabolic gene expression reveals clusters of cancer cell lines associated with tumor type and developmental lineage. Related to Figure 1. (A)** UMAP visualization of cell lines based on the expression levels of 1,620 metabolic genes. Cell lines are colored based on their cancer type. **(B)** Principal component analysis (PCA) of the expression of 1,620 metabolic genes in hematopoietic cancer cell lines. Cell lines are colored based on whether they belong to lymphoid or myeloid lineages (left) or based on cancer type (right). Variance explained by the first two principal components (PC) is shown. **(C)** Non-Small Cell Lung Cancer (NSCLC) cell lines and their subtypes, including Giant Cell Carcinoma of the Lung, Large Cell Lung Carcinoma, Lung Adenocarcinoma, Lung Adenosquamous Carcinoma, Lung Squamous Cell Carcinoma, Mucoepidermoid Carcinoma of the Lung, Non−Small Cell Lung Cancer, and Poorly Differentiated Non−Small Cell Lung Cancer, are highlighted on UMAP plot from Figure 1.

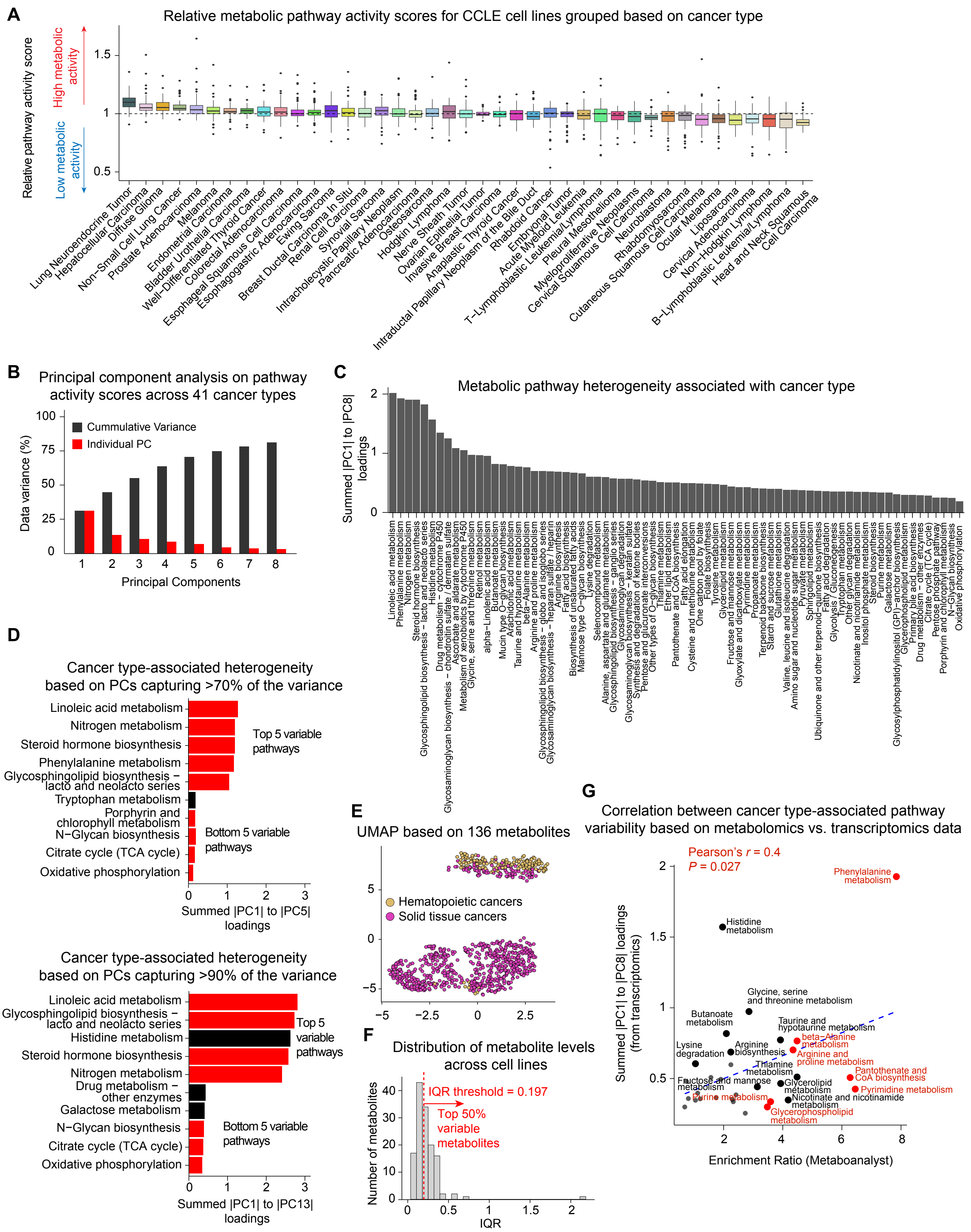


**Figure S2. Variability of the overall metabolic activity across cell lines of different cancer types. Related to Figure 2. (A)** Calculated metabolic activity scores (original, non-permutated scores) of 73 pathways across 41 cancer types. A metabolic pathway activity score < 1 for a cancer type represents reduced pathway activity in that cancer type in comparison with the average pathway activity across all cancer types; scores > 1 represent increased activity; and a score of 1 represents activity levels equivalent to the average over all cancer types. **(B)** Principal component analysis (PCA) of metabolic pathway activity scores across 41 cancer types. Percentage of variance captured by the first eight principal components (PC) are shown. **(C)** Ranking of metabolic pathways based on the extent to which their heterogeneity is associated with cancer type, evaluated by computing the absolute sum of PCA loadings for each pathway over the first eight principal components. **(D)** Ranking of metabolic pathways based on the extent to which their heterogeneity is associated with cancer type, evaluated by computing the absolute sum of PCA loadings for each pathway over the first five principal components capturing >70% of overall variance in data (top) or over the first thirteen principal components capturing >90% of overall variance. The top 5 and bottom 5 variable pathways are shown. Pathways that are found in the top/bottom 5 variable pathways based on 80% variance threshold (Figure 2B) are highlighted in red. **(E)** UMAP visualization of CCLE cell lines (n = 875 cell lines across 34 cancer types) based on the abundance of 136 metabolites. Cell lines are colored based on whether they associate with hematopoietic or solid tissue cancers. **(F)** Distribution of interquartile ranges (IQRs) for the median abundance of 136 metabolites in cell lines across 29 cancer types. Only cancer types with at least 5 cell lines profiled via metabolomics were included in the analysis. The tope 50% metabolites variable across cancer types were determined based on a 50-percentile IQR threshold (i.e., 0.197). **(G)** Correlation between variable pathways across cancer types identified through metabolomics vs. transcriptomics data analysis. Pathway enrichment analysis of the variable metabolites across cancer types was performed using metabolite set enrichment analysis (MSEA) based on KEGG database in Metaboanalyst. Enrichment ratio from Metaboanalyst was correlated against the absolute sum of PCA loadings for each pathway (**C**) computed based on transcriptomics data.

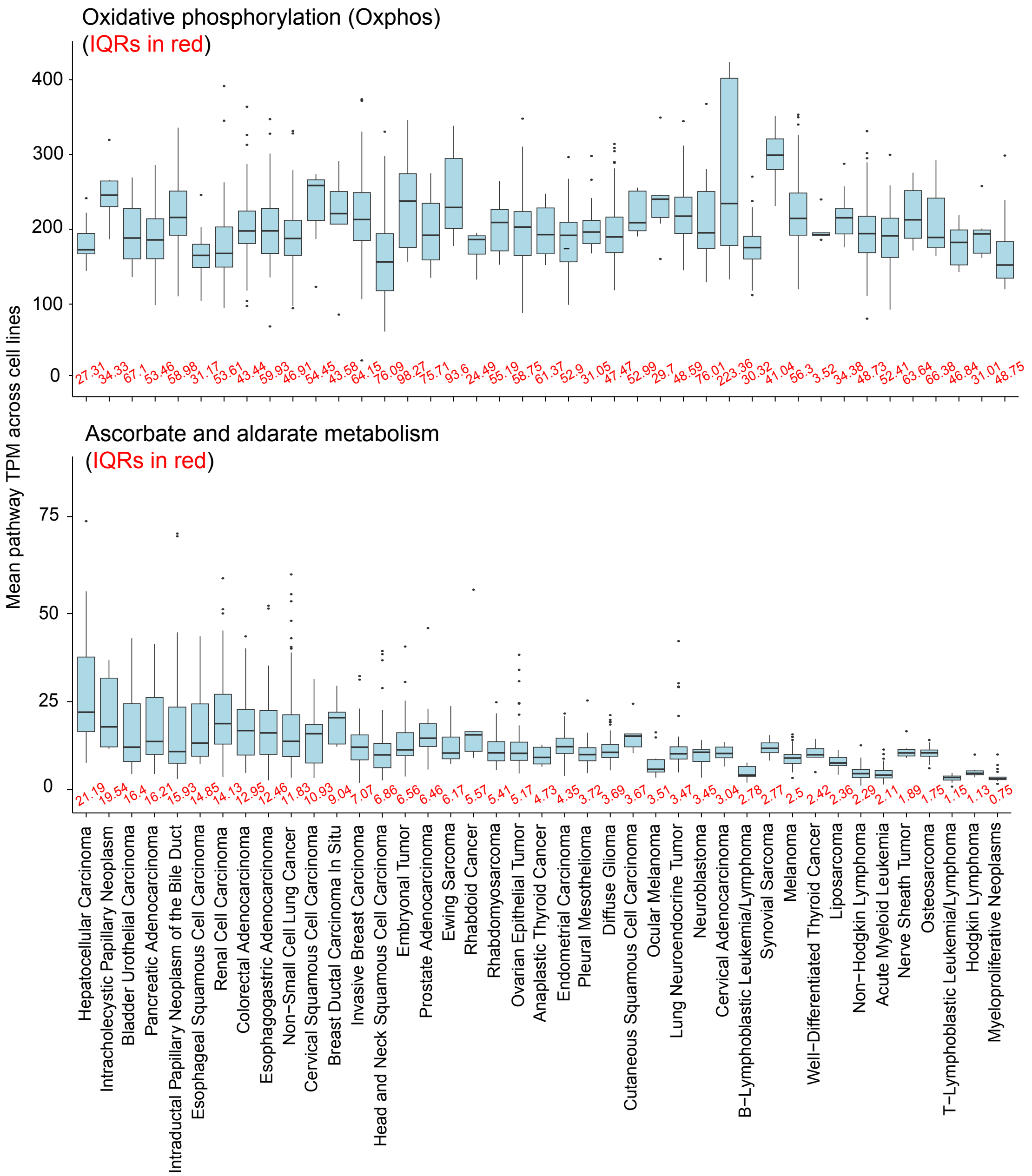


**Figure S3. Distinct patterns of cell line-to-cell line variability in two representative metabolic pathways. Related to Figure 3.** The data are shown for oxidative phosphorylation (top) and ascorbate and aldarate metabolism (bottom) across indicated cancer types. Mean pathway transcript per million (TPM) levels, their median and interquartile ranges (IQR) across cell lines are highlighted.

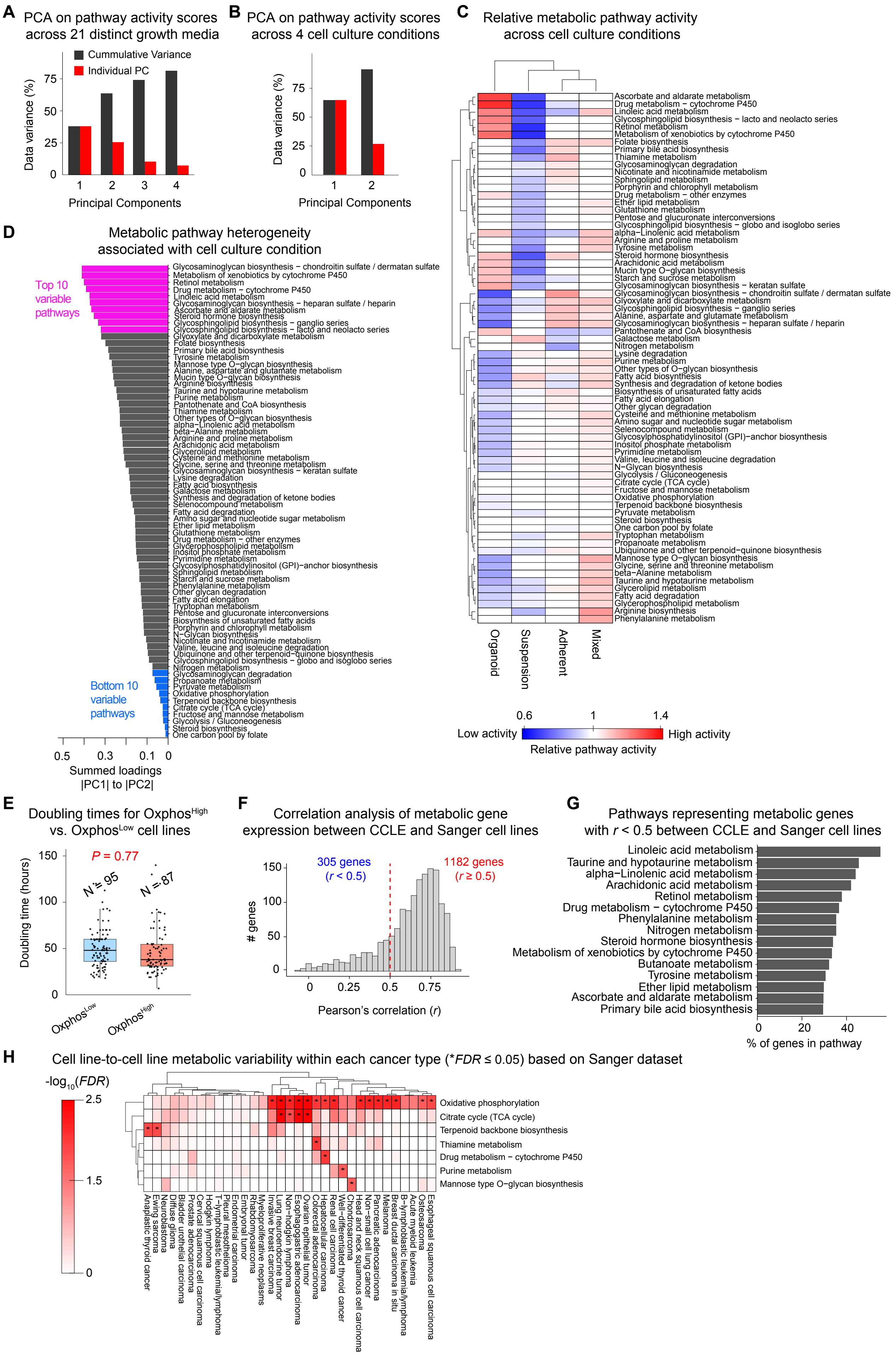

A
PCA on pathway activity scores across 21 distinct growth media
Cummulative Variance
Individual PC
Data variance (%)
Principal Components
B
PCA on pathway activity scores across 4 cell culture conditions
Data variance (%)
Principal Components
C
Relative metabolic pathway activity across cell culture conditions
Organoid
Suspension
Adherent
Mixed
Low activity
High activity
Relative pathway activity
Ascorbate and aldarate metabolism
Drug metabolism − cytochrome P450
Linoleic acid metabolism
Glycosphingolipid biosynthesis − lacto and neolacto series
Retinol metabolism
Metabolism of xenobiotics by cytochrome P450
Folate biosynthesis
Primary bile acid biosynthesis
Thiamine metabolism
Glycosaminoglycan degradation
Nicotinate and nicotinamide metabolism
Sphingolipid metabolism
Porphyrin and chlorophyll metabolism
Drug metabolism − other enzymes
Ether lipid metabolism
Glutathione metabolism
Pentose and glucuronate interconversions
Glycosphingolipid biosynthesis − globo and isoglobo series
alpha−Linolenic acid metabolism
Arginine and proline metabolism
Tyrosine metabolism
Steroid hormone biosynthesis
Arachidonic acid metabolism
Mucin type O−glycan biosynthesis
Starch and sucrose metabolism
Glycosaminoglycan biosynthesis − keratan sulfate
Glycosaminoglycan biosynthesis − chondroitin sulfate / dermatan sulfate
Glyoxylate and dicarboxylate metabolism
Glycosphingolipid biosynthesis − ganglio series
Alanine, aspartate and glutamate metabolism
Glycosaminoglycan biosynthesis − heparan sulfate / heparin
Pantothenate and CoA biosynthesis
Galactose metabolism
Nitrogen metabolism
Lysine degradation
Purine metabolism
Other types of O−glycan biosynthesis
Fatty acid biosynthesis
Synthesis and degradation of ketone bodies
Biosynthesis of unsaturated fatty acids
Fatty acid elongation
Other glycan degradation
Cysteine and methionine metabolism
Amino sugar and nucleotide sugar metabolism
Selenocompound metabolism
Glycosylphosphatidylinositol (GPI)−anchor biosynthesis
Inositol phosphate metabolism
Pyrimidine metabolism
Valine, leucine and isoleucine degradation
N−Glycan biosynthesis
Glycolysis / Gluconeogenesis
Citrate cycle (TCA cycle)
Fructose and mannose metabolism
Oxidative phosphorylation
Terpenoid backbone biosynthesis
Pyruvate metabolism
Steroid biosynthesis
One carbon pool by folate
Tryptophan metabolism
Propanoate metabolism
Ubiquinone and other terpenoid−quinone biosynthesis
Mannose type O−glycan biosynthesis
Glycine, serine and threonine metabolism
beta−Alanine metabolism
Taurine and hypotaurine metabolism
Glycerolipid metabolism
Fatty acid degradation
Glycerophospholipid metabolism
Arginine biosynthesis
Phenylalanine metabolism
D
Metabolic pathway heterogeneity associated with cell culture condition
Top 10 variable pathways
Bottom 10 variable pathways
Summed loadings |PC1| to |PC2|
E
Doubling times for Oxphos^High vs. Oxphos^Low cell lines
P = 0.77
N = 95
N = 87
Doubling time (hours)
Oxphos^Low
Oxphos^High
F
Correlation analysis of metabolic gene expression between CCLE and Sanger cell lines
305 genes (r < 0.5)
1182 genes (r ≥ 0.5)
### genes
Pearson's correlation (r)
G
Pathways representing metabolic genes with r < 0.5 between CCLE and Sanger cell lines
Linoleic acid metabolism
Taurine and hypotaurine metabolism
alpha−Linolenic acid metabolism
Arachidonic acid metabolism
Retinol metabolism
Drug metabolism − cytochrome P450
Phenylalanine metabolism
Nitrogen metabolism
Steroid hormone biosynthesis
Metabolism of xenobiotics by cytochrome P450
Butanoate metabolism
Tyrosine metabolism
Ether lipid metabolism
Ascorbate and aldarate metabolism
Primary bile acid biosynthesis
% of genes in pathway
H
Cell line-to-cell line metabolic variability within each cancer type (*FDR ≤ 0.05) based on Sanger dataset
-log10(FDR)
Oxidative phosphorylation
Citrate cycle (TCA cycle)
Terpenoid backbone biosynthesis
Thiamine metabolism
Drug metabolism − cytochrome P450
Purine metabolism
Mannose type O−glycan biosynthesis

**Figure S4. The impact of growth media, cell culture condition, doubling time (growth rate) on metabolic pathway activity of diverse cancer cell lines and their consistency across CCLE and Sanger datasets. Related to Figure 4. (A)** Principal component analysis (PCA) of metabolic pathway activity scores across 21 distinct growth media. Percentage of variance captured by the first four principal components (PC) capturing >80% of total variance in data are shown. **(B)** PCA of metabolic pathway activity scores across 4 distinct cell culture conditions. Percentage of variance captured by the first two PCs capturing >80% of total variance in data are shown. **(C)** Hierarchical clustering of relative activity scores for 73 significantly variable metabolic pathways across 4 cell culture conditions. A metabolic pathway activity score < 1 for a growth medium represents reduced pathway activity in that growth medium in comparison with the average pathway activity across all growth media; scores > 1 represent increased activity; and a score of 1 represents activity levels equivalent to the average over all growth media. **(D)** Ranking of metabolic pathways based on the extent to which their heterogeneity is associated with cell culture condition, evaluated by computing the absolute sum of PCA loadings for each pathway over the first two PCs. **(E)** Comparison of doubling time between $Oxphos^{High}$ and $Oxphos^{Low}$ cell lines. Statistical significance was evaluated by two-sided, two-sample *t* test. **(F)** Pearson's correlation analysis of metabolic gene expression across 1,487 metabolic genes and 658 cell lines shared between CCLE and Sanger datasets. **(G)** Pathways representing metabolic genes with Pearson's correlation coefficient of $r < 0.5$ between CCLE and Sanger cell lines. Metabolic pathways in which at least 30% of constituent genes had $r < 0.5$ are highlighted in order. **(H)** Metabolic pathways enriched in genes with highest contribution (determined by PCA) to the metabolic heterogeneities among individual Sanger cell lines from different cancer types. Data are clustered based on -$\log_{10}$(FDR) values derived from gene set enrichment analysis (GSEA). Pairs of pathway/cancer type with significant cell line-to-cell line heterogeneity ($FDR \leq 0.05$) are highlighted with an asterisk (*).

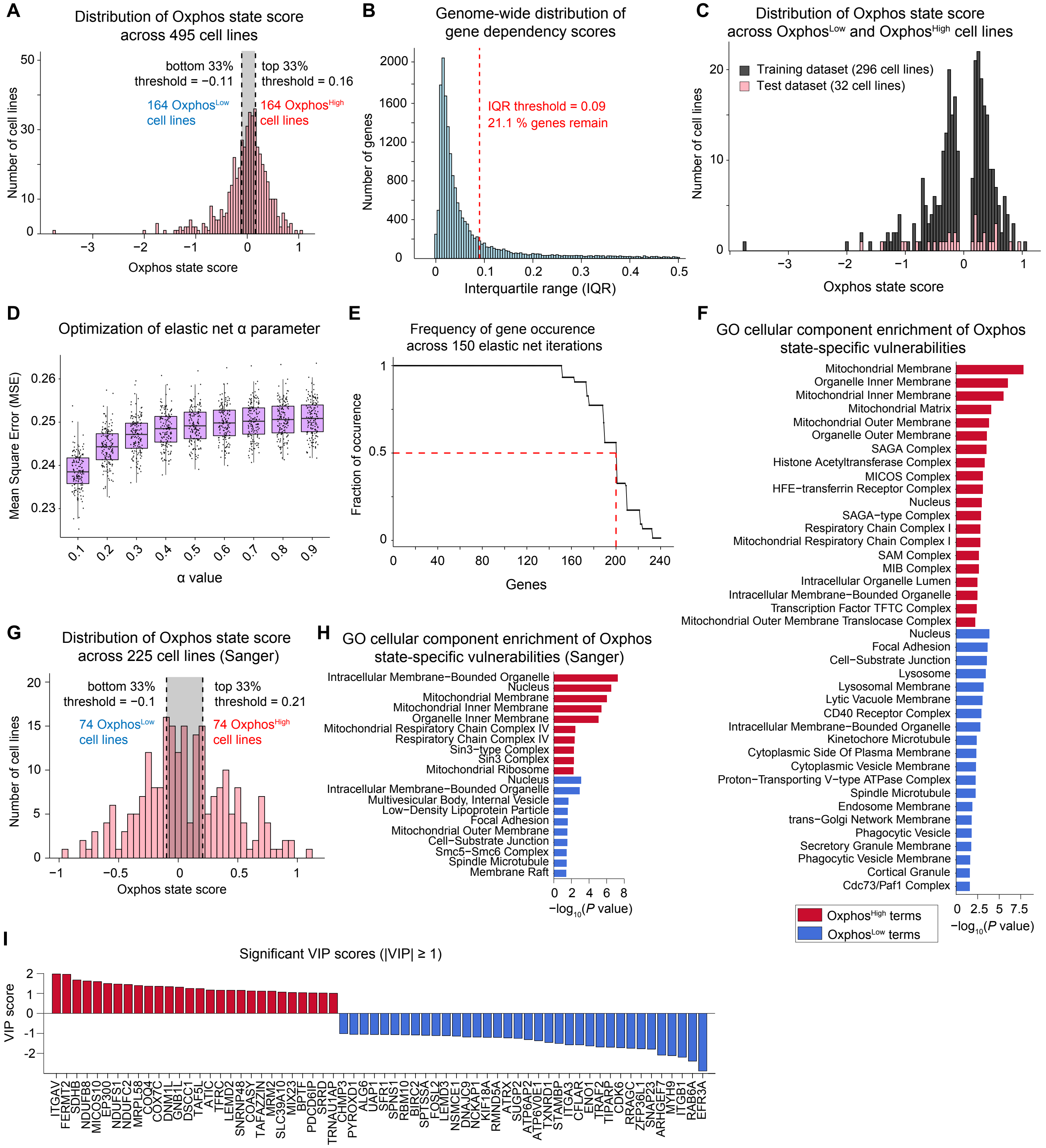


**Figure S5. Elastic net regularization selects a subset of gene features based on their association with the Oxphos state score across cell lines. Related to Figure 5. (A)** Distribution of oxidative phosphorylation (Oxphos) states scores across 495 cell lines. $Oxphos^{High}$ and $Oxphos^{Low}$ groups of cell lines (each containing 164 cell lines) were determined based on the top and bottom 33 percentiles. **(B)** Distribution of interquartile ranges (IQR) of gene dependency scores for 17,202 genes. IQR for each gene was calculated across 328 cell lines. IQR threshold (dotted red line) and remaining genes after IQR filtering (right of the dotted red line) are shown. **(C)** Distribution of Oxphos state scores across randomly selected training set (296 cell lines) and test set (32 cell lines). **(D)** Optimization of elastic net model parameter α. Boxplots of mean squared error (MSE) across 150 elastic net iterations (for each α) with varying α values of 0.1 to 0.9. **(E)** Fraction of occurrence for a given gene appearing across 150 random elastic net iterations. Genes that occurred in at least 50% of the iterations were used in the next step (i.e., PLSR modeling). **(F)** The top enriched GO cellular components associated with gene vulnerability features selected by elastic net regularization (*P* ≤ 0.05). Genes with positive and negative elastic net coefficients were used to infer vulnerabilities in $Oxphos^{High}$ and $Oxphos^{Low}$ cell lines, respectively. **(G)** Distribution of Oxphos state scores across 225 Sanger cell lines. $Oxphos^{High}$ and $Oxphos^{Low}$ groups of cell lines (each containing 74 cell lines) were determined based on the top and bottom 33 percentiles. **(H)** The top enriched GO cellular components associated with gene vulnerability features selected by elastic net regularization (*P* ≤ 0.05) using the Sanger dataset. Genes with positive and negative elastic net coefficients were used to infer vulnerabilities in $Oxphos^{High}$ and $Oxphos^{Low}$ cell lines, respectively. **(I)** PLSR-derived VIP scores ≥ 1 or ≤ -1. The sign of the VIP scores shows whether the identified gene is a vulnerability in $Oxphos^{High}$ cell lines (red) or $Oxphos^{Low}$ cell lines (blue).

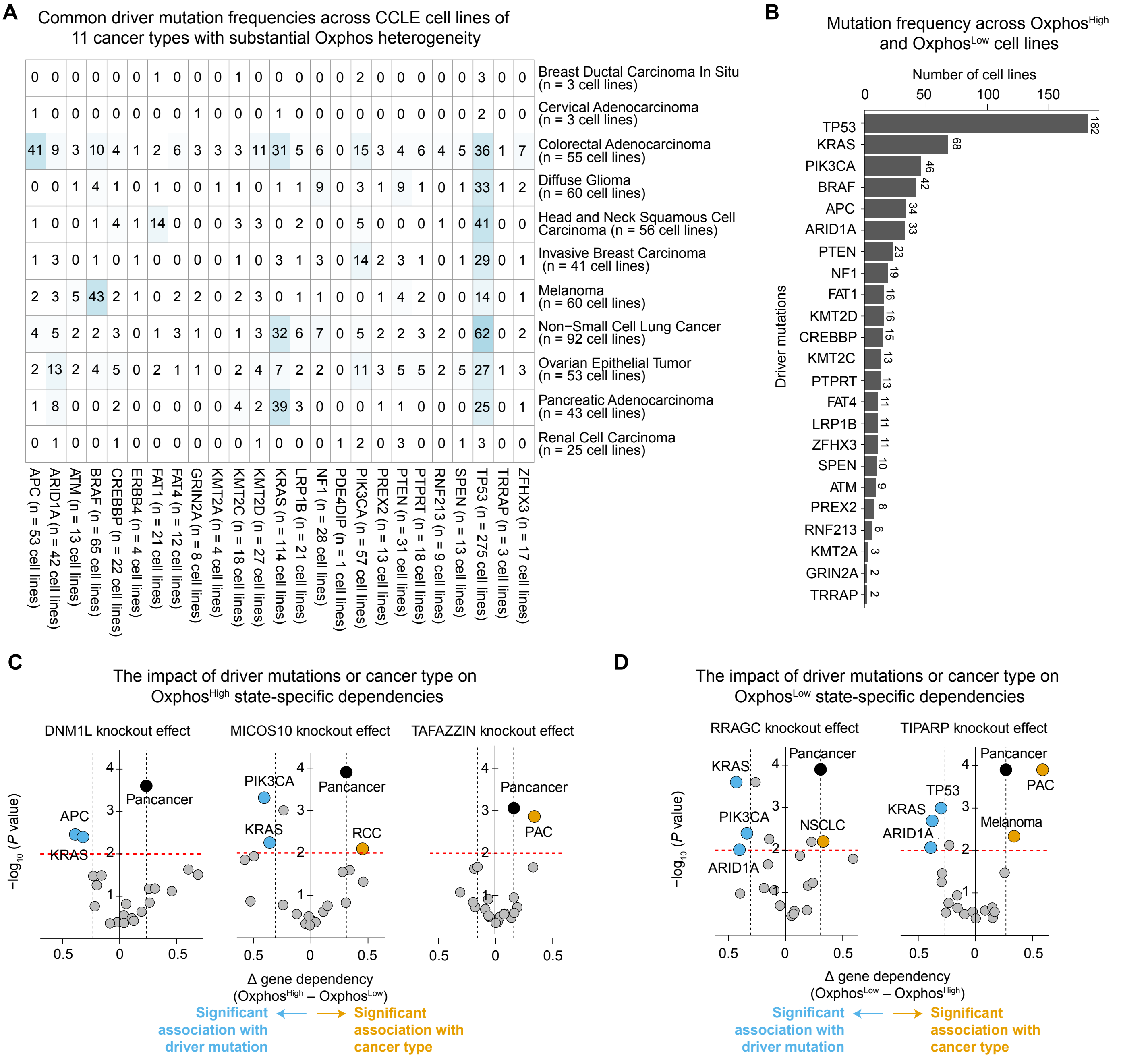


**Figure S6. Statistical analysis reveals synthetically lethal associations between Oxphos state, driver mutations and tissue context. Related to Figure 6. (A)** The number of cell lines bearing each of the top 25 most frequent driver mutations across eleven Oxphos-variable cancer types. Total mutation frequency for each of the 25 mutations is shown on the bottom. Total number of cell lines for each Oxphos-variable cancer type is shown on the right. **(B)** The top 25 most frequent driver mutations and their associated frequencies across $Oxphos^{High}$ and $Oxphos^{Low}$ cell lines. Number of cell lines that have a given mutation is shown next to each bar. **(C, D)** Statistical enrichment of $Oxphos^{High}$-associated **(C)** and $Oxphos^{low}$-associated **(D)** gene vulnerabilities in cell lines associated with specific tissue types or driver mutations. Enrichment is considered significant if the median gene dependency difference (Δ gene dependency) between $Oxphos^{High}$ and $Oxphos^{Low}$ subgroups of cell lines associated with a cancer type or driver mutation is significantly larger than the median gene dependency difference between $Oxphos^{High}$ and $Oxphos^{Low}$ groups regardless of cancer type and mutation (i.e., pan-cancer median difference level highlighted by the black circle and dotted line). Statistical significance of median differences was determined by empirical *P* values computed based on permutation testing ($P \leq 0.01$; red dotted line).

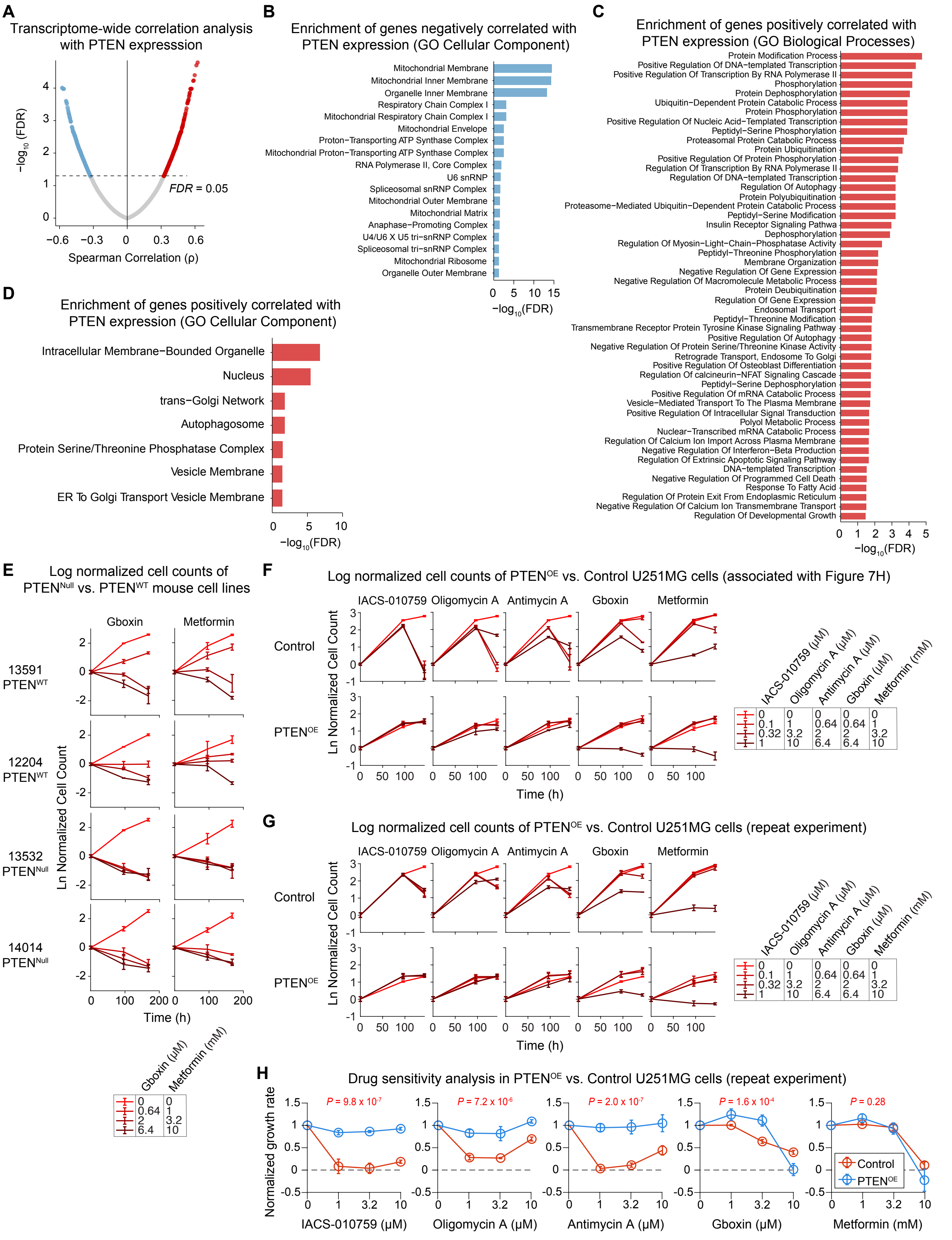


**Figure S7. Loss of PTEN predicts increased dependency on mitochondrial respiratory chain in Oxphos$^{High}$ tumor cells. Related to Figure 7. (A)** Spearman's correlation analysis to rank transcripts based on the association of their abundance with PTEN expression in 79 patient-derived glioma tumors**.** Transcripts negatively correlated (blue) or positively correlated (red)

with PTEN mRNA levels were highlighted as significant if $FDR \leq 0.05$. **(B)** The top Gene Ontology (GO) cellular components ($FDR \leq 0.05$) associated with genes whose expression negatively correlated with PTEN mRNA levels across 79 glioma tumors. **(C, D)** The top GO biological processes **(C)** and cellular components **(D)** associated with genes whose expression positively correlated with PTEN mRNA levels across 79 glioma tumors ($FDR \leq 0.05$). **(E)** Log-normalized changes in live cell count following exposure of $PTEN^{Null}$ (13532 and 14014) and $PTEN^{WT}$ (13591 and 12204) primary mouse glioma cell lines to Gboxin and metformin at indicated doses for a total period of 7 days. Data are presented as mean ± SD for across three replicates. **(F, G)** Log-normalized changes in live cell count following exposure of U251MG cells overexpressing PTEN ($PTEN^{OE}$) or Vector control (Control) to indicated concentrations of IACS-010759, oligomycin A, antimycin A, Gboxin, and metformin for a total period of 6 days. Data in **F** and **G** represent two independent experiments performed on different days using different batches of U251MG culture. **(H)** Normalized growth rates measured in U251MG cells overexpressing PTEN ($PTEN^{OE}$) or Vector control (Control) following treatment with indicated concentrations of IACS-010759, oligomycin A, antimycin A, Gboxin, and metformin using repeated experimental data shown in **G**. The average net growth rates were calculated from measurements of live cell count at three timepoints (including 0, 96, and 144 hours). Statistical significance of the effect of each drug on $PTEN^{OE}$ vs. Control cells was determined by two-way ANOVA, conducted specifically on the middle two doses of each drug (i.e., 0.1 and 0.32 μM for IACS-010759; 1 and 3.2 μM for oligomycin A; 0.64 and 2 μM for antimycin A and Gboxin; 1 and 3.2 mM for metformin). Data in **E-H** are presented as mean ± SD for across three replicates.
